# Overcoming sensory-memory interference in working memory circuits

**DOI:** 10.1101/2025.03.17.643652

**Authors:** Andrii Zahorodnii, Diego Mendoza-Halliday, Julio C. Martinez-Trujillo, Ning Qian, Robert Desimone, Christopher J. Cueva

## Abstract

Memories of recent stimuli are crucial for guiding behavior, but the sensory pathways responsible for encoding these memories are continuously bombarded by new sensory experiences. How the brain overcomes interference between sensory input and working memory representations remains largely unknown. To formalize the solution space, we examined recurrent neural networks that were either hand-designed or trained using gradient descent methods, and compared these models with neural data from two different macaque experiments. Here we report mechanisms by which neural networks overcome sensory-memory interference using both static and dynamic coding strategies: gating of the sensory inputs, modulating synapse strengths to achieve a strong attractor solution, and dynamic strategies – including the extreme solution in which cells invert their feature preference during working memory. Neural data from the medial superior temporal (MST) area of macaques, where sensory and working memory signals first interact along the dorsal pathway, best aligned with a solution that combined input gating and tuning inversion. Behavioral predictions from this model also matched error patterns observed in monkeys performing a working memory task with distractors. Taken together, our results help elucidate how working memory circuits preserve information as we continue to interact with the world, and suggest intermediate cortical visual areas like MST may play a critical role in this computation.

## 1 INTRODUCTION

Our behavior is guided both by immediate sensory experiences and by memories of recently encountered stimuli. Many studies explore how neural circuits store and maintain information in working memory (Baddeley and Hitch, 1974; Hopfield, 1982; Amit, 1989; Zipser, 1991; Miyake and Shah, 1999; Khona and Fiete, 2022). However, it remains largely unknown how the neural circuits that support working memory both allow information to flow into them while also preserving this information as we continue to interact with the world (Figure 1a) (Libby and Buschman, 2021; Cueva* et al., 2021). How do memory representations keep from being overwritten by new sensory inputs? This is the central question of this study.

**Figure 1.**
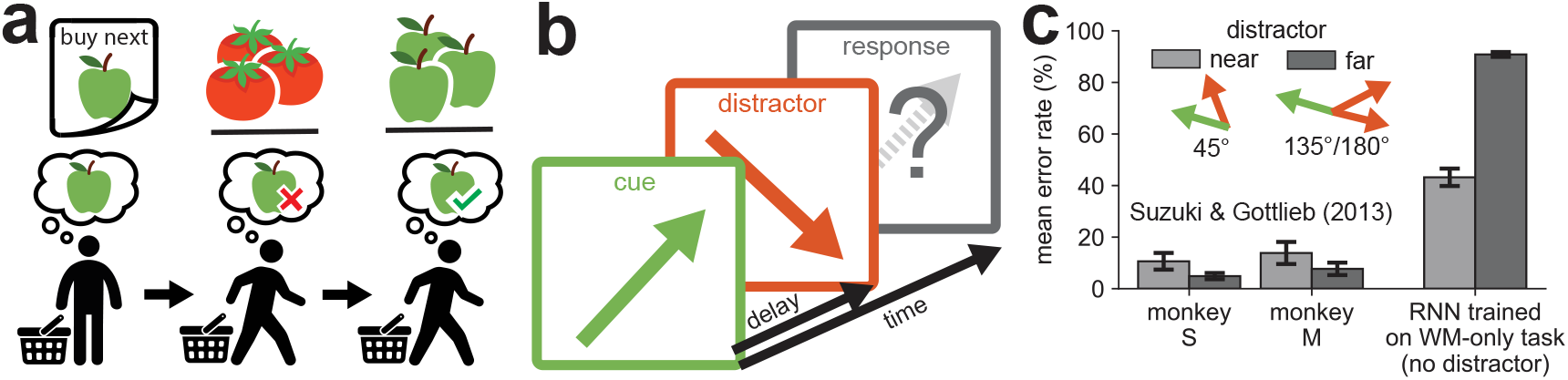
In recurrent neural networks, robustness to a distractor is an additional capability beyond working memory storage. (a) Sensory-memory interference in the real world, at the grocery store. We see an apple on our shopping list and then can keep this item in memory even as we see other items on our search through the store. (b) Working memory task with distractor. A cue is shown at one of eight directions uniformly spaced 45° apart. After a variable time interval, a distractor is shown in the WM+D condition. The goal of this task is to produce a response at the initial cue direction while ignoring the distractor. During the testing phase, the distractor can either be near the input stimulus (45° away) or far (135°/180° away). (c) Behavioral data (error rates) for two monkeys trained on the task. Adapted from Suzuki and Gottlieb (2013). Networks trained on the working memory only task without a distractor (WM-only) are not robust to the distractor (right). Error bars indicate s.e.m.

The problem of sensory-memory interference is likely widespread, as sensory features are often encoded by broadly tuned cells, for example, direction and orientation tuning in middle temporal (MT), and primary visual area (V1) neurons (Albright, 1984; Schiller et al., 1976). If two stimulus orientations are presented successively at the same location (Ding et al., 2017), they provide similar inputs, via the same set of connections, to the same set of memory units. How, then, does the system prevent the memory of the first orientation from being overwritten by the arrival of the second? Outside of controlled experimental settings, the problem of sensory-memory interference must still be overcome by neural circuits as eye movements realign relevant stimuli so we effectively have sequential presentations of stimuli in the same retinotopic positions, much like the experimental settings.

Here, we study the general problem of sensory-memory interference in the context of a class of problems that are common to many species, such as, humans (Ma et al., 2014), monkeys (Robertson et al., 1999; Pasternak and Greenlee, 2005; Wimmer et al., 2014), rats (Taube et al., 2000), mice (Yoder and Taube, 2009), flies (Seelig and V., 2015; Green et al., 2017; Kim et al., 2017), and fish (Petrucco et al., 2023); namely, remembering a circular variable that can take continuous values between 0 and 360 degrees. Theoretical models, like ring attractor circuits, have been proposed to remember these circular variables (Skaggs et al., 1994; Ben-Yishai et al., 1995; Redish et al., 1996; Zhang, 1996; Stringer et al., 2002; Xie et al., 2002). However, we show that a naive implementation of this memory system alters memories of recent stimuli with subsequent sensory inputs and does not agree with experimental results from monkey behavior.

To understand the underlying computations required to overcome the problem of sensory-memory interference, we trained, examined, and eventually were able to engineer, recurrent neural networks (RNNs) to solve this problem. We found an infinite RNN solution space that included gating of the sensory inputs, modulating synapse strengths to achieve a strong attractor solution, and dynamic visual feature coding, such that, at the extreme, cells invert their tuning over time. Each solution makes unique experimental predictions about 1) the connectivity between neurons, and 2) how neural tuning curves change over time. We tested whether the patterns of neuronal activity in our models were consistent with those observed during working memory tasks in two experiments (Mendoza-Halliday et al., 2014, 2024) with single-neuron electrophysiological recordings from cortical area medial superior temporal (MST). MST was identified as the first area along the dorsal visual pathway to show feature-selective sustained spiking activity during working memory in addition to selective visual responses (Mendoza-Halliday et al., 2014) and thus is of particular interest for our study on sensory-memory interference. Contrary to dominant theories of working memory, we found that in nearly half of the neurons in MST, the preferred motion direction during the working memory delay period was opposite to the preferred direction during the stimulus presentation period (Mendoza-Halliday et al., 2025). Interestingly, the neural activity in MST, which included these inversions of tuning, was well aligned with one of the models for overcoming sensory-memory interference. More specifically, the neural activity from MST was most aligned with the Gating + Inversion of Tuning solution. This solution was also consistent with experimental results from monkey behavior on a working memory task with distractors (Suzuki and Gottlieb, 2013). Taken together, our results show how recurrent neural networks are able to solve the problem of sensory-memory interference using a combination of both static and dynamic codes, and suggest that this solution may be implemented in intermediate visual processing stages such as area MST, where sensory and working memory signals first interact.

## 2 RESULTS

### 2.1 In recurrent neural networks, robustness to a distractor is an additional capability beyond working memory

To investigate the mechanisms by which a neural network might overcome the problem of sensory-memory interference, we study a working memory task with a distractor (Figure 1b). In this task, a cue stimulus with a particular feature value between 0 and 360 degrees is presented, followed by a delay period during which the cue is absent and must be remembered in order to subsequently report a correct response. At a variable time during the delay period, a task-irrelevant distractor stimulus with a different feature value is presented, and subjects must try to maintain the cue direction in working memory without interference by the distractor feature. The cue stimulus is encoded by a population of broadly tuned feature-selective cells (Teich and Qian, 2003), resembling a population of V1 or MT neurons with orientation or direction tuning curves (Schiller et al., 1976; Albright, 1984).

We trained continuous-time recurrent neural networks (RNNs) (Miller and Fumarola, 2012; Mante* et al., 2013) in two conditions: either on the full task with distractors (WM+D), or on the simple working memory only task (WM-only) where distractors were absent. After training, RNNs from both training conditions were able to solve the WM-only task (mean error *<*10°). However, RNNs in the WM-only training condition did not generalize to the distractor task. To compare these behavioral results with experimental data, we test the model on a variation of the WM+D task from Suzuki and Gottlieb (2013) where the cue and distractor are shown at one of eight directions uniformly spaced 45° apart. The distractor can either be similar (near) to the input stimulus (45° away) or far (135°/180° away) as shown in Figure 1b. In this task with discrete directions, the target selected by the RNN was taken to be the one nearest the RNN output. The RNNs trained without distractors exhibited error rates far greater than that of the monkeys in Suzuki and Gottlieb’s study (Figure 1c). Additionally, the RNNs had more errors for the far distractors than for the near distractors whereas the monkeys showed the opposite error pattern. From this finding, we concluded that in RNN models, the ability to withstand the distractor is dissociable from, and additional to, the ability to store a memory representation of the cue.

### 2.2 RNNs trained using gradient descent leverage both static connectivity and dynamic tuning to solve the task

To gain a mechanistic understanding of how trained RNNs solve the WM+D task, we examined their dynamics and connectivity (Figure 2).

**Figure 2.**
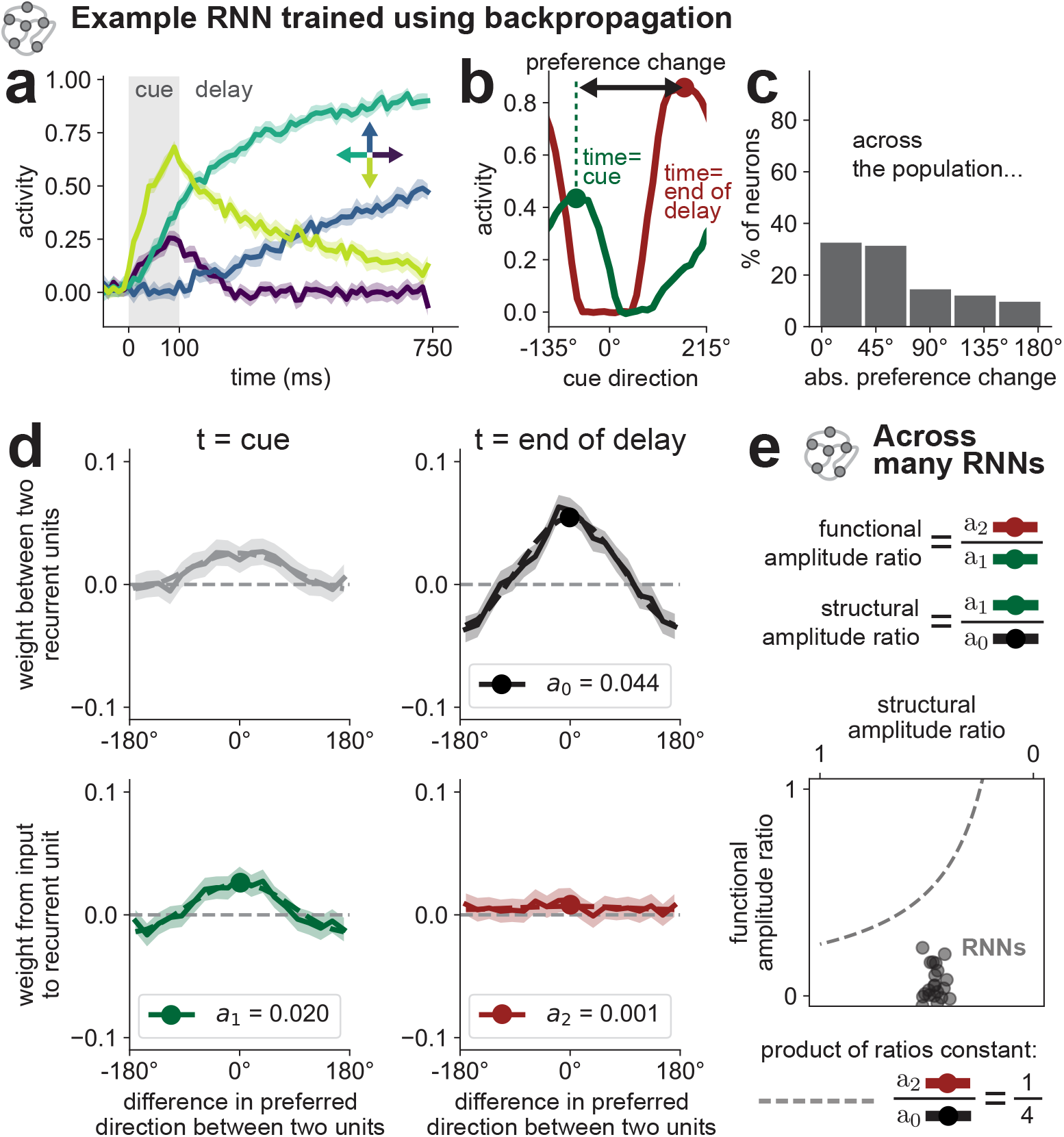
RNNs trained using gradient descent manipulate both structural and functional connectivity to solve the working memory plus distractor task. (a) Activity of one example artificial neuron (unit) when the cue stimulus is presented at four cardinal directions. Note the change in the unit’s preferred direction over time. (b) Tuning curves of the neuron from (a) at two points in time: in the middle of the cue period (t=cue), and at the end of the delay period (t=end of delay). Preference change is defined as the difference between the preferred directions at those two points in time. (c) Histogram showing the distribution of absolute preference changes of all direction-selective neurons in the example RNN. (d) Functional connectivity patterns between recurrent units (top row) and between the input and the recurrent units (bottom row) sampled at the two points in time (left and right columns). For every pair of units, the connection weight between them is plotted against the difference in their preferred cue directions at that point in time, and then all points are binned and averaged to produce a mean population curve. The shaded regions denote the 95% confidence interval of the mean. Cosine curves are fit to the connectivity patterns in all conditions, with amplitudes extracted (dashed lines; a=amplitude). (e) Scatter plot showing the structural and functional amplitude ratios across *N*=20 RNNs. Every point on the graph represents a backpropagation-trained network. The dotted line shows a constant product of structural and functional amplitude ratios, and indicates one set of networks with approximately equal performance (Supplementary Figure S3).

For any given collection of RNN units, we define the functional network connectivity pattern as the (binned and averaged) relationship between the weights connecting any pair of units, as a function of the difference in preferred stimulus directions for that pair of units. At the end of the delay period before the distractor, the pattern of functional network connectivity between direction-selective recurrent units resembled that of a ring attractor model with cosine connectivity (for an example network, see Figure 2d; top right). However, contrary to the prediction of the default ring attractor model, tuning curves of direction-selective units were not constant over time, and instead shifted largely between the cue presentation period and the end of the delay period (for an example unit, see Figure 2a, b). In other words, the recurrent units varied their preferred stimulus direction over time, instead of having a single fixed value (Cueva* et al., 2021). As a result of this transformation, most recurrent units changed their preferred input stimulus direction, as evidenced by a wide distribution of preference changes in the population (Figure 2c).

The functional impact of these tuning changes is to decrease the effective strength of the input onto the memory representation. The quantify the effect of this transformation, we looked at the functional network connectivity between input direction-selective units and the recurrent units before and after the transformation (Figure 2d, bottom). The amplitude of the connectivity pattern dramatically decreased as a result of the transformation; both patterns were fit well by cosine curves with amplitudes decreasing from *a*_1_ ==0.02 to *a*_2_ =0.001 for the example RNN. Note that the effective decrease in connection strength from the input to the recurrent units that occurs from the time of the cue period to the delay period (left and right columns in Figure 2d), are only driven by tuning changes in the population (weights are always kept constant after training). This finding suggested that the function of the transformation may be to make the effect of the future distractor input weaker and more diffuse, and thus decrease its impact on the memory representation.

Furthermore, note that because of this transformation, the amplitude of the excitatory connectivity pattern between recurrent units increases from the time of the cue to the end of the delay period (when the working memory is stored; *a*_0_ =0.044; Figure 2d, top right), and it is larger than that of the input projection’s excitatory connectivity at the time of the cue (when stimulus information enters the network; *a*_1_ =0.02; Figure 2d, bottom left).

We observed similar results across all trained RNNs (Figure 2e). We define the functional amplitude ratio as *a*_2_*/a*_1_, and the structural amplitude ratio as *a*_1_*/a*_0_ (in the notation above). The functional amplitude ratio quantifies how much the tuning changes of units in the network serve to increase or decrease the impact of the input stimulus at the time of the cue versus at the end of the delay period. The structural amplitude ratio is a measure of the strength of the input versus recurrent connectivity, and can be thought of as quantifying the impact of the stimulus relative to the memory attractor, or as the “strength of the attractor.” Across RNNs, we found the functional amplitude ratio =0.062 *±*0.080, and the structural amplitude ratio =0.463 *±*0.036 (mean*±* s.d., *N* =20 networks). In the next section we will show that achieving a low value for either of the two ratios is sufficient to solve the task.

### 2.3 Hand-designed RNNs perform comparably to trained networks, while solving the task using distinct strategies

There are multiple ways of solving the WM+D task in RNNs. In backpropagation-trained networks, the solution strategies are mixed; in addition, other strategies have been described in the literature that are not present in the trained RNNs (for example, gating). This prompted us to consider building RNNs by hand to only implement one solution at a time.

Here, we propose four hand-designed (engineered) RNNs that solve the task using four distinct solution strategies: Gating, Strong Attractor, Reshuffle of Tuning, and Inversion of Tuning (Figure 3). All solutions are based on the ring attractor architecture, which has a long history of experimental support at the level of neural activity, and is also architecturally consistent with regions in both invertebrate and vertebrate brains (Kim et al., 2017; Hulse et al., 2021; Petrucco et al., 2023). We note that all of our hand-designed RNNs are able to solve the distractor task, with performance comparable to that of backpropagation-trained networks (Figure 3i).

**Figure 3.**
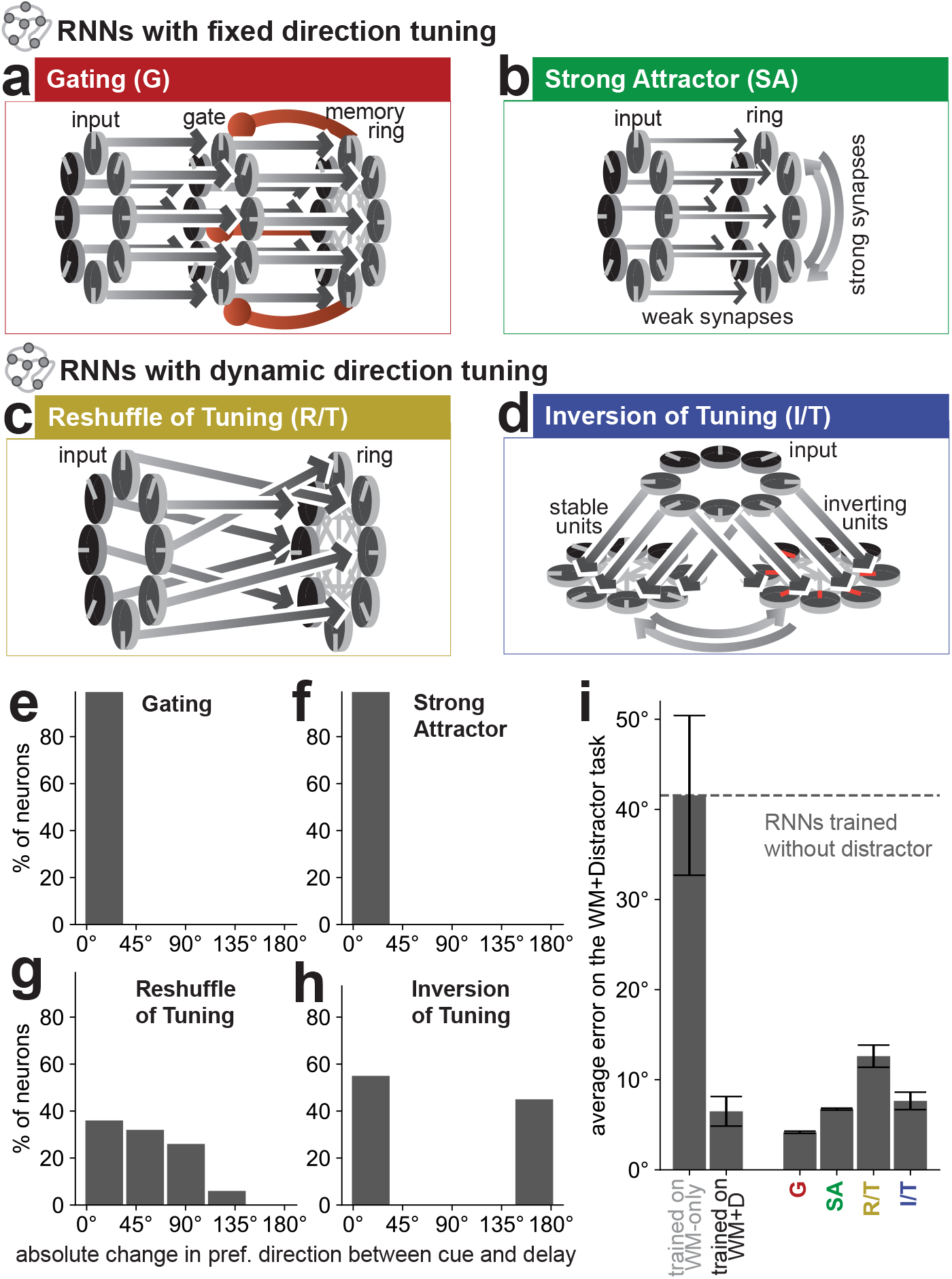
Hand-designed RNNs perform comparably to backpropagation-trained networks, while solving the task using distinct strategies. Each RNN makes distinct experimental predictions about the connectivity between neurons, and how the preferred directions change between the time of the initial cue and the subsequent delay period. (a-d) Hand-designed RNNs that solve the task using four distinct solution strategies: Gating (G; a), Strong Attractor (SA; b), Reshuffle of Tuning (R/T; c), Inversion of Tuning (I/T; d). All rings are based on the ring attractor connectivity. (e-h) Histograms showing predictions of each model for the distribution of preference changes of the neurons. (i) Performance of backpropagation-trained as well as hand-designed networks on the distractor task. Y-axis represents the mean angular error between the output of the network and the desired output across many trials. Error bars indicate the standard deviation across networks initiated from different random seeds. *N*=30 networks for every bar, with networks that did not converge omitted.

By hand-designing RNNs to solve the WM+D task, we accomplish two goals: (1) provide interpretable models that give conceptual understanding of the mechanisms that may be at play to solve the sensory-memory interference problem in trained RNNs and in biological networks and (2) confirm that the mechanisms described in this study can be sufficient on their own to overcome sensory-memory interference.

#### 2.3.1 Gating (G)

In the Gating solution (Figures 3a, 3e), information about the input stimulus flows through a cluster of units (a gate), which retranslates the same information to the memory cluster (a ring attractor). The gate is inhibited by the activity of the memory ring, and this inhibition protects the first sensory input.

#### 2.3.2 Strong Attractor (SA)

In the Strong Attractor solution (Figures 3b, 3f), the overall amplitude of the recurrent weights is much larger than that of the input connection weights, achieving a low structural amplitude ratio (*a*_1_*/a*_0_ *<* 0.25). After information about the first input stimulus (the cue) enters the network, the recurrent activity quickly ramps up to dominate the inputs to each of the recurrent units. After that point, the distractor is not able to substantially alter the stored information because the input projection is comparatively weak.

#### 2.3.3 Reshuffle of Tuning (R/T)

The Reshuffle of Tuning solution achieves a low functional amplitude ratio (*a*_2_*/a*_1_ *<* 0.25) by introducing misalignment between the input projections and the recurrent connectivity (Figure 3c; for details of the mechanism, see Supplementary Figure S1). For every recurrent unit, its misalignment with the input projection is chosen randomly and independently from a uniform distribution centered at 0°. In other words, each unit in the RNN has both a preferred direction for the input stimulus, determined by the input-to-recurrent connectivity, and a preferred direction for the memory representation, determined by the connectivity among the recurrent units; and these two preferred directions are not the same. Therefore, units change their tuning curves over time in the delay period immediately following the first input (Figure 3g), similar to the backpropagation-trained networks. These changing tuning curves, triggered by the initial input stimulus and guided by the patterns of connectivity, make the effect of the distractor more diffuse and thus decrease its impact on the stored memory.

#### 2.3.4 Inversion of Tuning (I/T)

The Inversion of Tuning solution, similarly to the Reshuffle of Tuning, achieves a low functional amplitude ratio (*a*_2_*/a*_1_ *≈*0.1) through a misalignment of input projections. However, in this solution misalignments are not picked randomly from a uniform distribution. Instead, two subclusters of units are defined: stable and inverting units (Figure 3d). The stable units receive aligned projections as normal. The inverting units receive projections that are misaligned by exactly 180°, i.e. receive strong drive from direction-selective inputs that have opposite tuning to each units’ preferred direction in the ring attractor. The stable and inverting units are connected together as a single ring attractor with the cosine connectivity pattern. Stable units slightly outnumber inverting units, which leads to dynamics where the stable units keep their tuning throughout the task, and the inverting units shift their tuning curves by 180° (invert their tuning; Figure 3h). The introduced competition between the stable and inverting units weakens the input projections relative to the recurrent weights, which decreases the impact of the distractor.

### 2.4 RNNs can solve the task while interpolating between solution mechanisms

The impact of sensory stimuli on the memory representation can be modulated by manipulating either the structural amplitude ratio (by decreasing the strength of input connections relative to the recurrent weights) or the functional amplitude ratio (by introducing misalignment between input and recurrent connectivity), or both (Figure 4d). In fact, we were able to engineer Strong Attractor + Reshuffle of Tuning (SA+R/T) RNNs that can solve the task by relying on either of those strategies to any predefined degree (Figure 4a). As long as the combined effect of the two mechanisms (quantified as the product of the structural and functional amplitude ratios, *a*_2_*/a*_0_) was low enough (in our experiments, *a*_2_*/a*_0_ *<* 0.25), the network performance was high on the WM+D task. The product of functional and structural amplitude ratios strongly predicted the performance of the hand-designed network (Supplementary Figure S3). However, a low overall ratio was not required for high performance on the WM-only task (Figure 4b). As I/T and R/T are based on the same mechanism, the same finding holds for SA+I/T networks (Figure 4c). As such, we found that those two strategies can be dissociated in networks, and are independent of each other, but can be combined in any given network.

**Figure 4.**
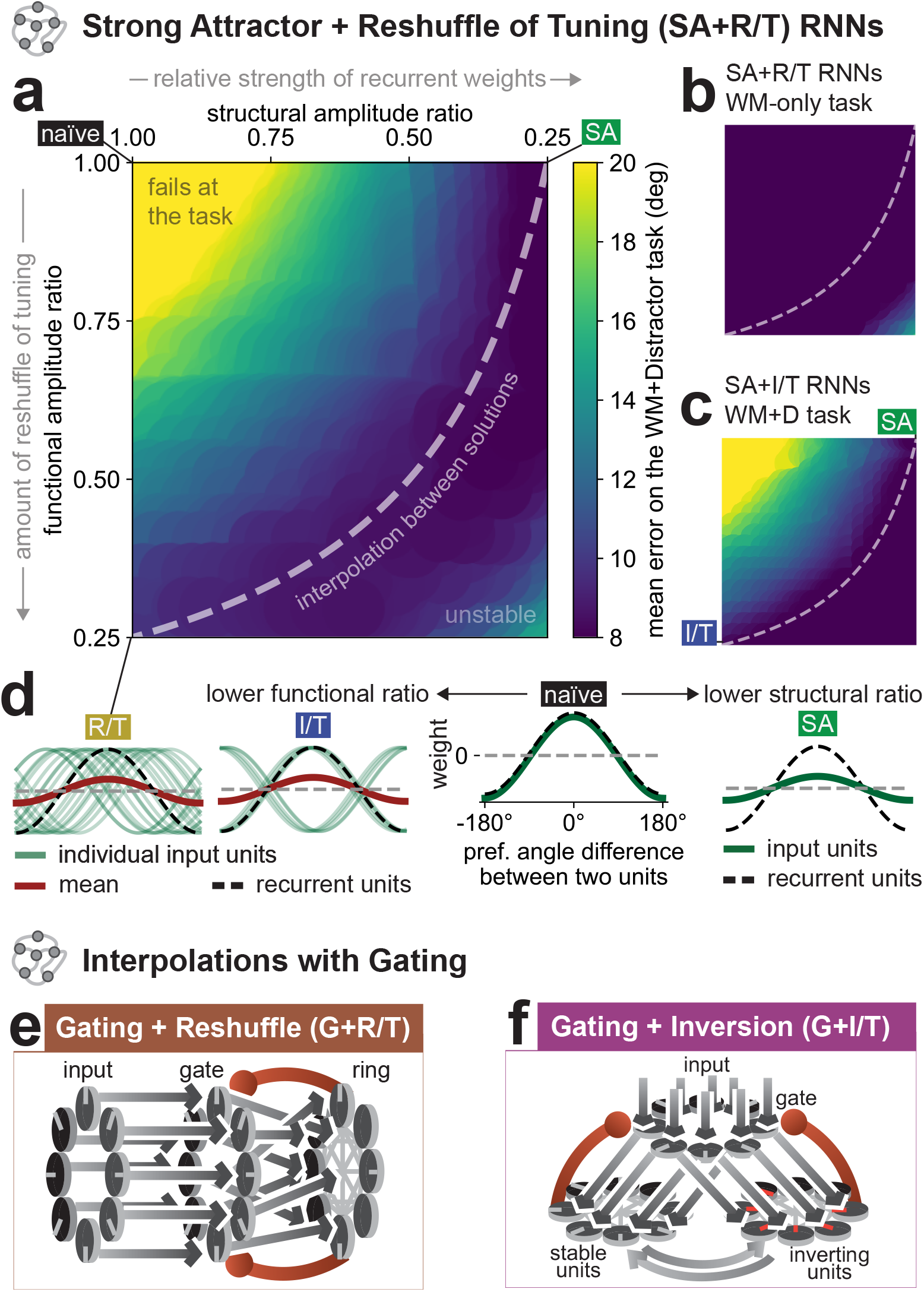
RNNs can solve the task while interpolating between solution mechanisms. (a) Heatmap of mean errors on the WM+D task across the different choices of structural and functional amplitude ratios. Every point on the heatmap represents a hand-designed Strong Attractor + Reshuffle of Tuning RNN, where the strength of recurrent connectivity and amount of functional tuning changes are varied independently. *N* =6639 networks. (b) Corresponding to (a) heatmap for the working memory only task, without the distractor. *N* =6639 networks. (c) Corresponding to (a) heatmap for *N* =5501 Strong Attractor + Inversion of Tuning RNNs on the WM+D task. (d) Diagram depicting the mechanisms by which different solution strategies impact the functional and structural ratios. (e, f) Connectivity of hand-designed Gating + Reshuffle of Tuning (G+R/T; e) and Gating + Inversion of Tuning (G+I/T; f) solutions.

In addition, by adding a layer of gate units, any solution can be combined with the Gating solution (Figures 4e, 4f; G+SA not shown). Finally, by manipulating the distribution of input-recurrent curve misalignments, interpolations can be obtained between the Reshuffle of Tuning solution and the Inversion of Tuning solution (data not shown).

Thus, interpolations can be obtained between any two solution strategies introduced in this study. In conclusion, we show that there exists an infinite solution space for the problem of sensory-memory interference, with behavioral performance that can be organized according to structural and functional amplitude ratios to quantify solution mechanisms that leverage both static connectivity and dynamic tuning.

### 2.5 Different solution mechanisms have distinct neural signatures

To inform future analyses, we looked for ways of experimentally differentiating between the solution mechanisms. We found that in the delay period immediately following the input stimulus, our models made distinct predictions for the distribution of preference changes in the population of direction selective neurons. Neurons in the Gating and Strong Attractor solutions maintained their preference throughout the task (Figures 3e, 3f). In the Reshuffle of Tuning, and Gating + Reshuffle of Tuning solution, we found a diffuse pattern of preference changes that spanned the full range of possible values (Figure 3g). Finally, the Inversion of Tuning, and Gating + Inversion of Tuning RNNs had two clusters of neurons: those that maintained their preference, and those that inverted to the opposite direction (Figure 3h).

These findings allowed us to narrow our hypothesis space when analyzing neural data and artificial networks. For example, our backpropagation-trained RNNs exhibited preference changes which were most consistent with the Reshuffle of Tuning mechanism (compare Figure 2c and Figure 3g).

### 2.6 Neurons in macaque brain area MST exhibit inversion of tuning, consistent with the Inversion of Tuning and Gating+Inversion of Tuning RNN solutions

To investigate how the brain might overcome sensory-memory interference, we examined neural data from two previously reported macaque experiments by Mendoza-Halliday et al. 2014 and 2024 (Mendoza-Halliday et al., 2014, 2024). In Mendoza-Halliday et al. 2014 monkeys performed a delayed match-to-sample (DMS) task, and in Mendoza-Halliday et al. 2024 monkeys performed a working memory-guided feature attention task. Both experiments requires (as a subproblem) overcoming sensory-memory interference, as some trials contained distractor stimuli before the behavioral response. The first experiment used spatially local random dot motion stimuli and the second used full-screen random dot motion stimuli. For a more complete description of the tasks, refer to the original studies (Mendoza-Halliday et al., 2014, 2024). In the present study, we looked specifically into the neural dynamics during the presentation of the first stimulus and the immediately following delay period.

We focused on neural responses in the brain area medial superior temporal. The findings of Mendoza-Halliday et al. (2014) show that MST has working memory activity, and suggest that this is the first area along the cortical visual processing stream where sensory processing signals encounter working memory signals (and thus is of interest in our study). Additionally, MST may play a substantial role in cognitive computation, for example in encoding abstract categorical decisions (Wild and Treue, 2021; Zhou et al., 2022).

Results from both studies showed that in MST, there is a combination of neurons that preserve their motion direction preference, and neurons that invert their preference, during the delay period with respect to the cue period (for example neurons, see Figures 5a, 5c). In both cases, the distributions of absolute preference changes contained two peaks, at 0° and 180° (Figures 5b, 5d), most consistent with the Inversion of Tuning mechanism, as well as the Gating + Inversion of Tuning mechanism.

**Figure 5.**
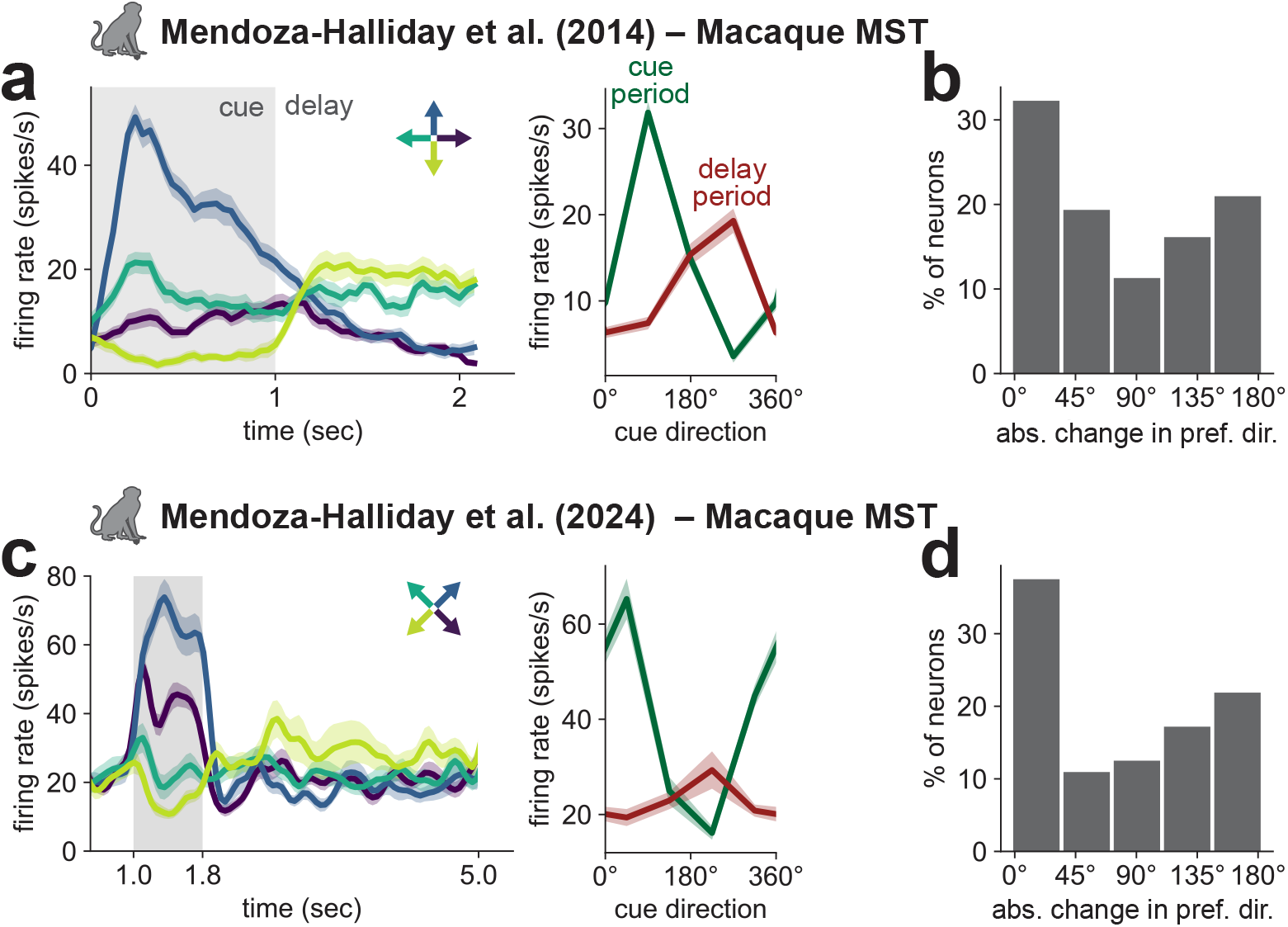
Neurons from the macaque brain area MST exhibit inversion of tuning properties, consistent with the Inversion of Tuning and Gating + Inversion of Tuning RNN solutions. (a-d) Neural data from macaque brain area medial superior temporal (MST) in two experiments: (a) Mendoza-Halliday et al. 2014 and (c) Mendoza-Halliday et al. 2024. Left, mean firing rate across trials during the cue and delay periods for an example MST neuron. Right, the tuning curves in the two time windows for the corresponding neurons. (b, d) Histogram of changes in preferred direction (absolute value) between the cue and delay period across all selective MST neurons recorded in the 2014 (b) and 2024 (d) studies.

Are these inverting feature preferences due to neural adaptation? If so, then we would predict that the inverting neurons would lose their direction selectivity during the delay period (Benda, 2021; Van Wezel and Britten, 2002; Bisley et al., 2004; Glasser et al., 2011; Murray et al., 2014). However, this is not what we observed. The direction selectivity of the inverting neurons is not monotonically decreasing during the delay, and in fact, even increases between the first and second half of the 3200 ms delay period in the experiment of Mendoza-Halliday et al. 2024 (Supplementary Figure S2, *p <* 0.01, Wilcoxon signed-rank test). It is possible that the timescale of decay for neural adaptation in MST is so large that we could not detect it across the 3200 ms delay period. However, as we show in section 2.8 the noise correlations from simultaneously recorded pairs of MST neurons are consistent with the delay period activity being maintained by ring attractor dynamics, and so we do not think that neural adaptation is *maintaining* the inverting feature preferences during the delay.

Are these inverting feature preferences *initiated* by neural adaptation? Neural adaptation is a complex phenomenon arising from the interplay between intrinsic neuronal properties and local circuit dynamics, including interactions among distinct cell types within cortical microcircuits. These mechanisms may contribute to the emergence of the inverted tuning observed in MST (Koch et al., 2025). Despite this potential complexity, our models offer a sufficient mechanism for these inversions of tuning. In our models we know precisely what is driving this phenomena, namely, each inverting neuron has both a preferred direction for the input stimulus, determined by the input-to-recurrent connectivity, and a preferred direction for the memory representation, determined by the connectivity among the recurrent units; and these two preferred directions are misaligned by 180 degrees. This misalignment causes neurons to invert their feature preference. In light of our results with artificial networks, our experimental findings suggest that the dynamic neural coding of the sensory representation that takes place in MST after the disappearance of the stimulus may be to protect the newly obtained memory representation from potential sensory interference.

### 2.7 Procrustes distance analysis suggests that neural responses in MST are most aligned with the Gating + Inversion of Tuning mechanism

To more directly and quantitatively compare the neural responses to those generated by our introduced solution strategies, we compared the pattern of activity from the entire population of recorded neurons to the pattern of activity from all units in each model. We used the Procrustes distance metric (Williams et al., 2021; Ding et al., 2021), which can be viewed as the residual distance after two neural trajectories are aligned with an optimal rotation (Harvey et al., 2023). Using this metric, we computed pairwise distances between all models and neural recordings, using cue-period and delay-period firing rates from every neuron (Supplementary Figure S4).

Using t-SNE, a nonlinear dimensionality reduction technique (van der Maaten and Hinton, 2008), we embedded all models and neural data as points in a 2d space - the solution space for the sensory-memory interference problem (Figure 6a). We found that by embedding the points in this low-dimensional space, the clustering of networks by solution type was recovered.

**Figure 6.**
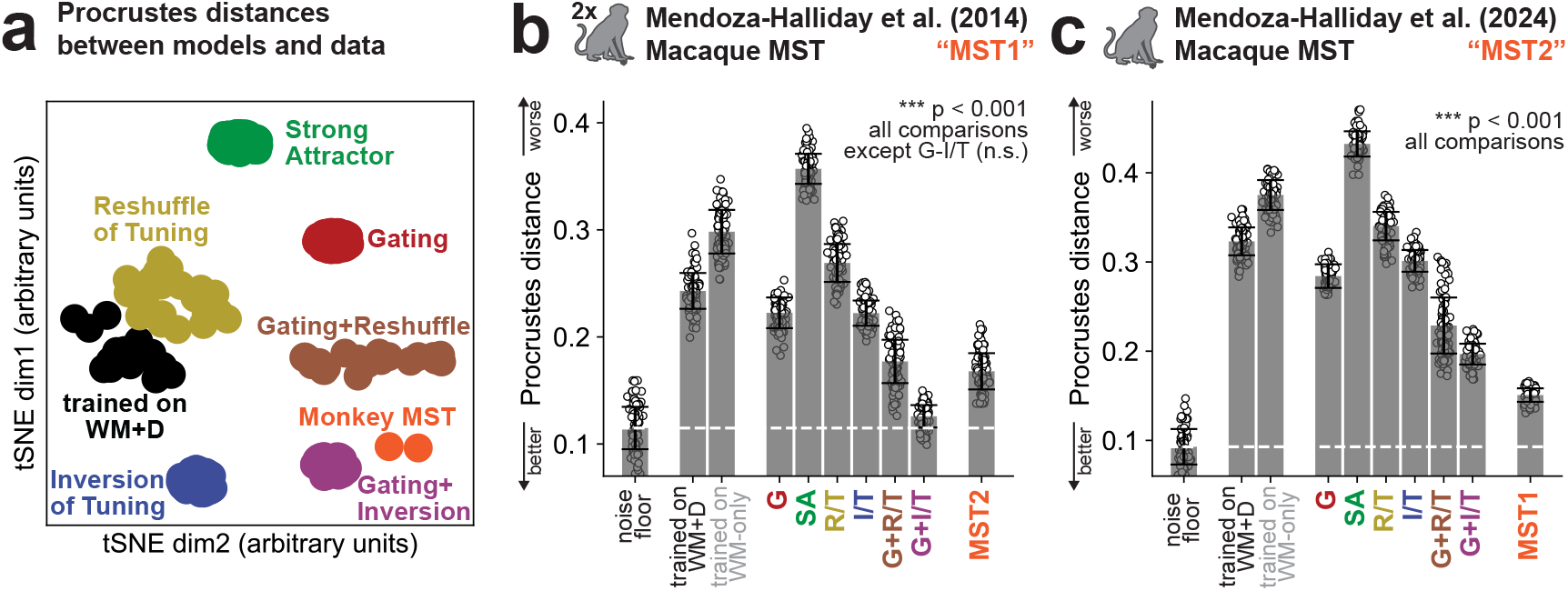
Neural responses from the medial superior temporal area of macaques are most aligned with the Gating + Inversion of Tuning mechanism. (a) The solution space of models and data is visualized using the t-SNE algorithm on the firing rates of neurons in the cue and delay periods. Every point represents a model network or neural dataset. (b, c) Each datapoint is the Procrustes distance between the firing rates from all units in a model and neurons from macaque brain area MST (Mendoza-Halliday et al., 2014, 2024). Across both experiments, neural data from MST is most aligned with the Gating + Inversion of Tuning mechanism. Error bars indicate the standard deviation. For (b), *p <* 0.001 for all comparisons except G+I/T (n.s.); for (c), *p <* 0.001 for all comparisons.

We found that data from both macaque experiments had the smallest distance to the G+I/T solution (0.13 ± 0.01 and 0.20 ± 0.01 for the 2014 and 2024 experiments, respectively, as shown in Figures 6b and 6c). In fact, in the experiments of Mendoza-Halliday et al. 2014, the distance to the G+I/T model (0.13 ± 0.01) was not significantly different from the noise floor (0.11 ± 0.02). The observed Procrustes distances were, overall, reflective of the similarity of the distribution of feature preference changes between each RNN (Figures 3e-h) and the experimental datasets (Figures 5b,d). Notably, the lowest Procrustes distances occurred for RNNs that included inversions of feature preference, a property observed in both datasets. This finding was robust irrespective of whether we included neurons selective for the cue direction in either the cue *or* delay periods (Figures 6b and 6c), or in both the cue *and* delay periods (Supplementary Figure S5).

### 2.8 Noise correlations in MST depend on the cue direction as predicted by the ring attractor model

The similarity between the neural activity in MST and the G+I/T model suggests that the underlying circuit in MST, responsible for the sustained activity during the delay period, is governed by ring attractor dynamics. This hypothesis of ring attractor dynamics makes a strong experimental prediction, namely, that noise correlations between pairs of simultaneously recorded neurons depend on the cue direction, as predicted by the ring attractor model (Ben-Yishai et al., 1995; Pouget et al., 1998; Wu et al., 2008, 2016). Even when the cue direction is the same across trials, the bump location along the ring attractor may have small variations, leading neural activity to vary characteristically due to the recurrent connectivity between neurons. Specifically, in a ring attractor, if the cue direction is between the preferred directions of two neurons (in-flank condition; Figure 7a), then a shift in the activity bump across trials will lead to increased activity in one neuron and decreased activity in the other; the activity will be negatively correlated across trials for repeated presentations of the same cue direction (Ponce-Alvarez et al., 2013; Wimmer et al., 2014). Intuitively, this is because the cue excites the two neurons at points along their tuning curves with opposite slopes, and fluctuations along these slopes are tied together across neurons due to coherent shifts in the bump activity. When the cue is shifted and presented outside the preferred directions of the two neurons, where both tuning curves have slopes of the same sign, the noise correlations are predicted to be positive (out-flank condition; Figure 7a). For values of the cue intermediate between these two extremes, near the peak of either of the two neurons’ preferred directions, the average noise correlations are predicted to be near zero. In Figure 7 we define the peak condition as occurring when the cue is within a 15 degree window centered around either neuron’s preferred direction.

**Figure 7.**
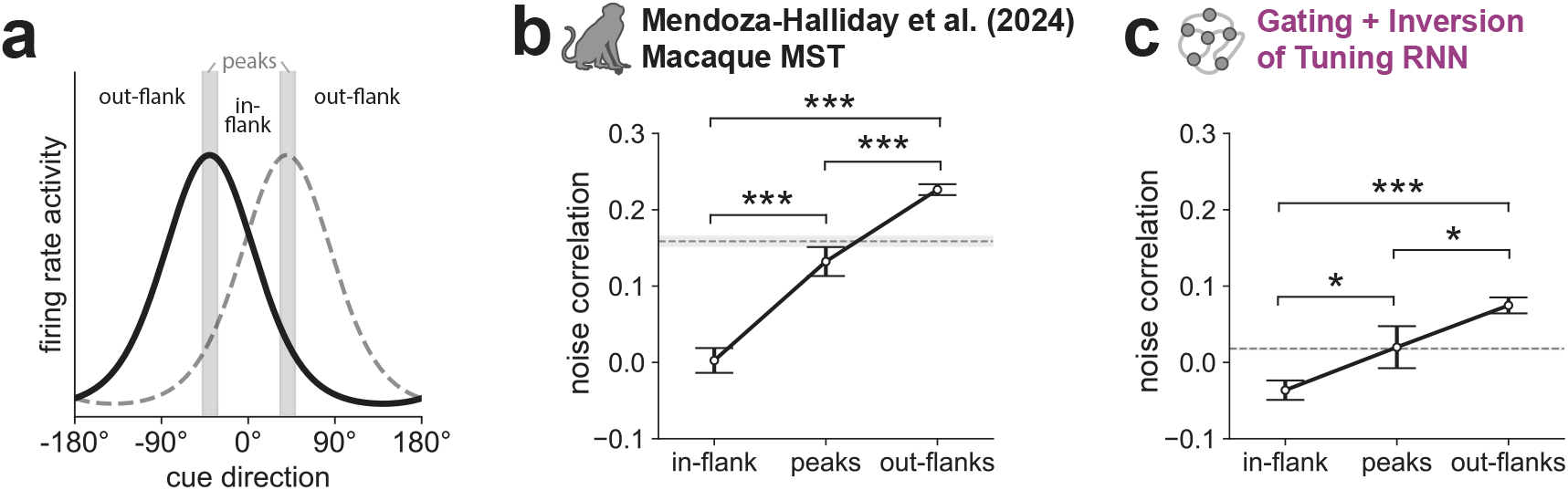
Noise correlations between pairs of neurons vary as the cue direction is changed, in agreement with the ring attractor model. (a) When the cue direction is in the in-flank, peak, or out-flank regions of the two tuning curves, the noise correlations between these two neurons are predicted to be ordered if they are part of a ring attractor circuit: in-flank *<* peaks *<* out-flanks. (b, c) Noise correlations during the delay period are ordered with in-flank *<* peaks *<* out-flanks for both MST and the G+I/T model (\**p <* 0.05, \*\*\**p <* 0.001, one-sided *t*-test). The horizontal shaded regions indicate the mean value during the fixation period before cue onset (baseline). In-flank correlations are less than baseline for both the MST data and G+I/T model (*p <* 0.001, one-sided *t*-test), while out-flanks correlations are greater than baseline (*p <* 0.001, one-sided *t*-test). Error bars indicate s.e.m.

These patterns of noise correlations are a signature of ring attractor dynamics but note that they can be uniformly increased by adding correlated inputs to the neurons (Hennequin et al., 2018) and indeed this offset is what we see across the pairs of simultaneously recorded neurons from Mendoza-Halliday et al. 2024 (*N* =225, 311, and 792 comparisons between neuron pairs for the in-flank, peaks, and out-flanks conditions, respectively; Figure 7b). We computed noise correlations during the 3.2 second delay period (excluding the 200 ms transient immediately following cue offset) and found that they were ordered with in-flank *<* peaks *<* out-flanks as predicted by the ring attractor model (Figures 7b,c; *p <* 0.001 for MST and *p <* 0.05 for the model, one-sided *t*-test). Additionally, if we take the average noise correlation during the fixation period before cue onset as a baseline (horizontal shaded region in Figures 7b,c) then we observe that this baseline is higher in MST than in the G+I/T model (consistent with correlated inputs in MST but not in the model) and that in-flank correlations are less than baseline for both the MST data and G+I/T model (*p <* 0.001, one-sided *t*-test), while the out-flanks correlations are greater than baseline (*p <* 0.001, one-sided *t*-test). In summary, noise correlations changed with cue direction as predicted by the ring attractor model.

### 2.9 Behavioral predictions of Gating + Inversion of Tuning solution, but not of pure Inversion of Tuning, are aligned with experimental behavioral data

In addition to making predictions about the neuronal activity patterns, the RNN models also make predictions about behavioral performance in a working memory task with distractors, and more specifically, about behavior as a function of the proximity of the distractor feature to the cue feature. Because the experimental design in the studies of Mendoza-Halliday et al. 2014 and 2024 did not allow us to thoroughly investigate the effects of distractors, we looked for other tasks to test behavioral predictions. We compared our models with behavioral responses from Suzuki and Gottlieb (2013) in which monkeys remembered the location of a cue stimulus while a distractor stimulus was presented at a different location during the delay period. This allowed us to analyze behavioral performance as a function of the similarity between cue and distractor features.

Models trained on the working memory task without a distractor, and the Inversion of Tuning models predict that the error rates should be higher when the distractor stimulus is in the direction opposite or near opposite to the cue stimulus (Figures 8b and 8c), which is not consistent with monkey behavioral data from Suzuki and Gottlieb (2013) (Figure 8a). However, the addition of Gating to the Inversion of Tuning solution changed the behavioral predictions of the model to be consistent with the experimental results (Figure 8d). This result reinforced the importance of the Gating component in the Gating + Inversion of Tuning solution as crucial for aligning the model with experimental data.

**Figure 8.**
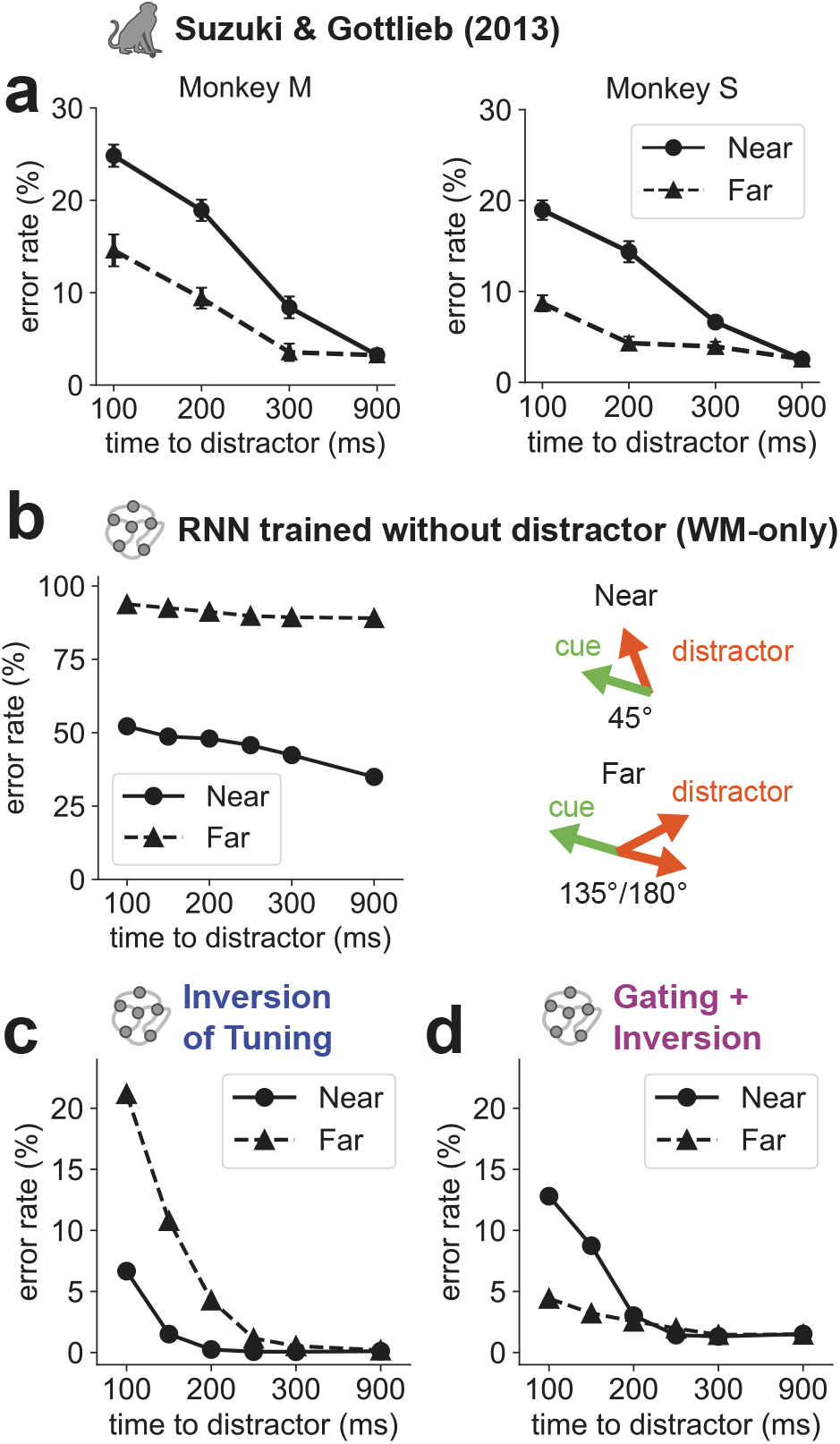
Addition of Gating to the Inversion of Tuning solution aligns behavioral predictions of the model with experimental data. (a) Behavioral data (error rates) on the working memory plus distractor task for two monkeys from Suzuki and Gottlieb (2013). The distractor can either be similar (near) to the initial cue stimulus (45° away) or far (135°*/*180° away). (b) A standard ring attractor network is not robust to the distractor and strongly alters memories of recent stimuli with subsequent inputs. (c-d) Behavioral patterns for the Inversion of Tuning (c) and Gating+Inversion of Tuning (d) hand-designed RNN solutions, corresponding to the behavioral experiment of Suzuki and Gottlieb (2013).

What’s driving these patterns of errors in the models? The behavior of the model is determined by the location of the activity bump. If this activity bump is centered over the cue location then the behavioral readout will be correct. If this activity bump is shifted far enough from the cue location by the distractor then the model response will be closer to one of the non-cue directions resulting in an error. This shift can happen because units with preferred directions that are not at the cue direction (referred to as “side units” in Figure 9) increase their activity. More specifically, errors occur when units with preferred directions on one side of the cue direction become more active after the distractor stimulus is presented, and units on the other side become less active (Figure 9a); we refer to this as the “differential modulation of side units”. This differential modulation of side units is shown in Figure 9b and well captures the behavioral errors for the models shown in Figure 8.

**Figure 9.**
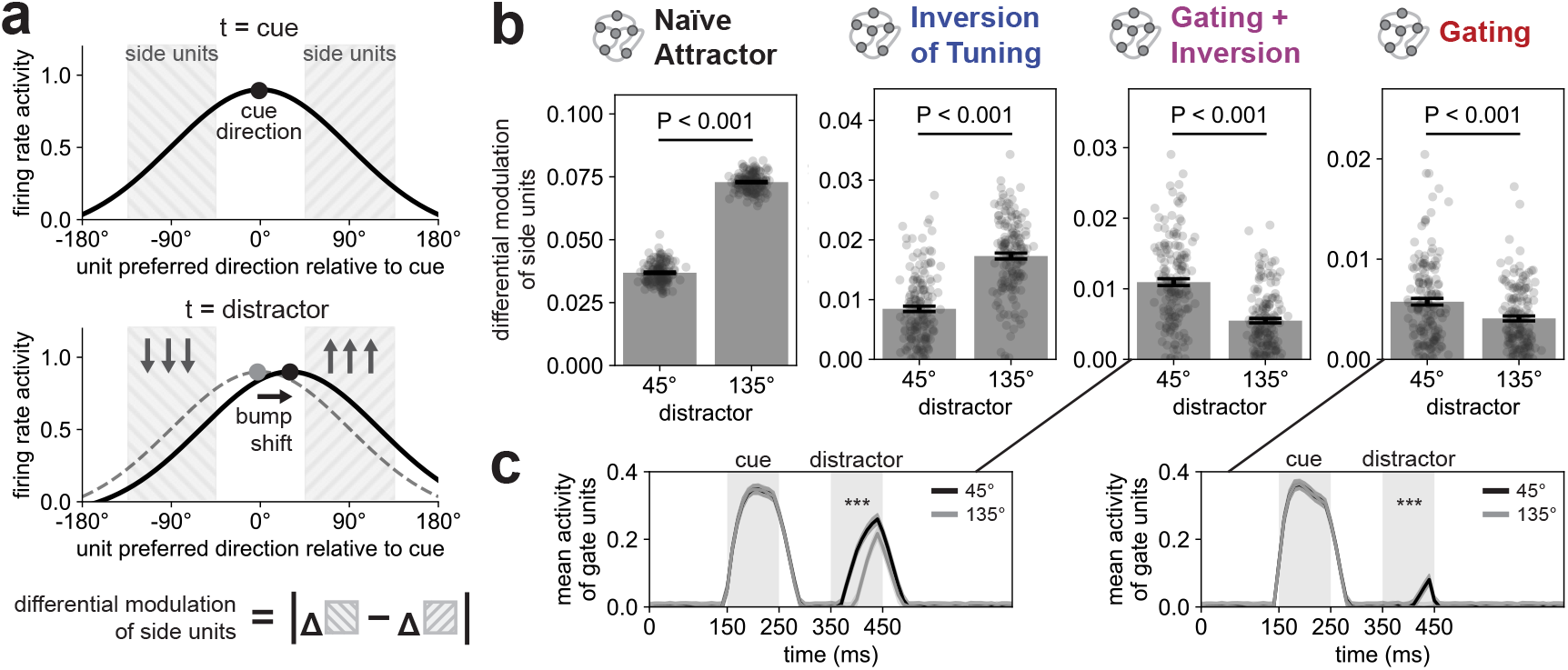
In RNNs, differential modulation of side units explains the patterns of behavioral error rates. (a) The differential modulation of side units is calculated as the absolute difference in changes of firing rates on the two flanks of the bump attractor. Higher values capture a shift of the bump from its original location during the distractor period, indicating a likely behavioral error. (b) Comparison of differential modulation of side units for an example near (45°) and far (135°) distractor position relative to the cue for various types of engineered RNNs: Naive Attractor, Inversion of Tuning, Gating, and Gating+Inversion of Tuning. This analysis reveals that for Naive Attractor and Inversion of Tuning, a distractor farther away induces a higher differential modulation of side units, however for the solutions including gating this pattern is reversed. *p <* 0.001, all comparisons. (c) The increased activity of gate units in the near distractor condition (for solutions involving gating) explains the higher differential modulation of side units. Error bars indicate s.e.m. \*\*\**p <* 0.001.

For example, the standard naive attractor has qualitatively similar error patterns as the WM-only model with more errors for distractors that are 135 degrees versus 45 degrees away from the cue direction, and correspondingly, for both models the differential modulation of side units is larger for distractors that are 135 degrees versus 45 degrees away from the cue direction. Both of these models do not know how to “ignore” a distractor, and a far distractor shifts the activity bump more than the near distractor does. Similarly, the Inversion of Tuning network has more errors for distractors that are 135 degrees versus 45 degrees away from the cue direction, and correspondingly, the differential modulation of side units is larger for distractors that are 135 degrees versus 45 degrees away from the cue direction. More intuitively, when the cue stimulus enters the Inversion of Tuning network it starts a competition between two bumps: the one generated by the stable units and the one generated by the inverting units. The stable units win this competition when no distractor is presented. However, if a distractor is presented far from the cue direction then this will add strength to the inverting bump, altering the balance of the competition and producing an error.

The networks with gating produce the opposite pattern of behavioral errors. There are more errors, and the differential modulation of side units is larger, when distractors are 45 versus 135 degrees away from the cue direction. Intuitively, the reason the near distractor achieves a higher error rate is that it can preferentially pass through the gate because those gate units were recently active (because they were transmitting the nearby cue) and inhibition from the ring has not yet closed the gate (Figure 9c).

An important caveat with the behavioral comparison in Figure 8 is that the study of Suzuki and Gottlieb (2013) used visuo-spatial locations around a fixation point rather than a variable along a single continuous circular dimension such as motion direction. Though there are differences between these two types of tasks, there is also experimental evidence that the underlying neural mechanisms rely on continuous ring attractors in both cases. Indeed, working memory for visuo-spatial locations appears to be governed by continuous ring attractor dynamics (Wimmer et al., 2014). In addition, by demonstrating the close similarity between our ring attractor model and MST data (Figures 6b, 6c, and Figure 7) our results suggest that the underlying circuit in MST, responsible for the sustained activity during the delay, is compatible with a ring attractor where both inverting and stably-tuned neurons are part of the same computational circuit. We further tested the match between MST data and predictions from such an integrated circuit model across all cell types by verifying that the noise correlations between pairs of stable-stable, inverting-inverting, and stable-inverting neurons are similarly high for similarly tuned neurons while correlations across all types of neuron pairs decrease as the difference in preferred stimulus grows (Supplementary Figures S6 and S7) (Leavitt et al., 2017b).

More generally, the circuit mechanisms we propose for overcoming sensory-memory interference are theoretically beneficial for ring attractor networks across different areas and modalities, and we hypothesize that they are likely employed beyond MST. There is indeed evidence that these inversions of tuning are also present in primary auditory cortex to reduce interference between sensory and memory representations (Libby and Buschman, 2021). In conclusion, we present the behavioral results for our models alongside the results from Suzuki and Gottlieb (2013) both as a suggestive parallel between behavioral outcomes when working memory circuits are governed by ring attractor dynamics, and to illustrate predictions to test by introducing a new distractor design in the tasks from Mendoza-Halliday et al. 2014 and 2024. Our results therefore not only demonstrate the presence of inversion of tuning in MST, but also propose a model circuit implementation protecting memories against interference that is consistent with an array of experimental results and leads to testable hypotheses.

## 3 DISCUSSION

Many studies explore how neural circuits store and maintain information in working memory. However, these circuits are not isolated memory storage devices. Instead, they must collaborate with sensory areas that feed information into them. Thinking about working memory in this larger context immediately raises new questions. How do the neural circuits that support working memory allow information to flow into them while also preserving this information from being overwritten under the constant bombardment of new sensory inputs as we continue to interact with the world? How does the brain overcome interference between sensory inputs and memory representations? Our insights are twofold – both in a computational and in a neurophysiological framework. In the computational framework, we show how RNNs can overcome sensory-memory interference by combining both static and dynamic neural codes that can be traded off to maintain constant error (Figure 4a). This understanding of the solution space allowed us to engineer solutions that could not be constructed before, and that were not found by standard task-optimization of RNNs. In the neurophysiological framework, we show that in macaque monkeys, after a stimulus is presented, the distribution of direction preferences in brain area medial superior temporal becomes bimodal, with nearly half of the neurons completely inverting their tuning. These responses challenge dominant theories of working memory in which a neuron would be expected to maintain its tuning preference from the cue period through the delay period. The neural activity in MST is most consistent with the Gating + Inversion of Tuning solution for overcoming sensory-memory interference.

It was particularly interesting to us that one of the RNN mechanisms that can serve to overcome sensory-memory interference - the inversion of tuning mechanism - was found to be a property of neurons in macaque area MST. Why would this mechanisms be observed particularly in area MST? One likely reason is that MST is the first cortical area along the dorsal visual pathway to show sustained spiking activity encoding motion directions in working memory (Mendoza-Halliday et al., 2014). In contrast, in cortical area MT, immediately upstream, neurons encode stimulus directions only during sensory input, but not when the stimulus is removed and remembered. Given that the transformation of sensory representations into mnemonic representations seems to occur in MST (Mendoza-Halliday et al., 2014), it is reasonable to expect that mechanisms to overcome sensory-memory interference and to maintain sensory and memory representations separate and disambiguated would be most important in this area, as well as in other intermediate-level sensory processing areas of the brain where sensory and memory signals first interact.

Our results indicate that the cortical architecture in MST, which allow neurons to generate sustained activity during working memory, is governed by ring attractor dynamics (Figures 6b and 6c, and Figure 7), and that both inverting and stably-tuned neurons are part of the same computational circuit. Consistent with this, the noise correlations between pairs of stable-stable, inverting-inverting, and stable-inverting neurons are similarly high for similarly tuned neurons and decrease as the difference in preferred stimulus grows (Supplementary Figures S6 and S7).

Libby and Buschman (2021), in the context of a flexible evidence integration task, also discovered neurons which inverted their feature preference, and argued for their significance in protecting the memory representation. Which mechanisms generate the stable and switching dynamics? Our study provides one answer to this question: the stable and switching dynamics in the Inversion of Tuning solution (which corresponds to the structured rotation mechanism of Libby and Buschman) arise as a result of the recurrent interactions within the network, and structured misalignments with the input projections. One implication of our results is that these dynamics do not require feedback projections from downstream areas, like prefrontal cortex, and can emerge as a result of local connectivity within a brain area. However, it is important to point out that behavioral goals - such as the decision about which sensory stimuli to maintain in working memory and which not to - are likely carried out by higher-order areas in parietal and prefrontal cortex rather than in MST. Therefore, it is possible that the mechanisms that protect memory representations from sensory interference also involve feedback from higher-order areas. Moreover, the idea that protection from sensory-memory interference acts at the level of one single brain area or population of neurons such as MST rests on the assumption that delay activity in that particular area plays a causal role in working memory storage. However, it could also be possible that delay period activity in MST is not causally involved in storage but serves a different purpose. Instead, storage mechanisms, and therefore mechanisms to protect from sensory-memory interference and to maintain segregation between sensory and memory coding, could be carried out in a different cortical area downstream, such as the lateral prefrontal cortex (Mendoza-Halliday and Martinez-Trujillo, 2017) or in a more widely distributed network of other areas (Leavitt et al., 2017a; Roussy et al., 2021).

There are multiple forces that guide any given network (artificial or biological) to a specific solution mechanism. When considering biological organisms, the diversity of possible factors can be broadly divided into two categories: nature (e.g. evolutionary factors) and nurture (e.g. training curriculum). It is possible that evolutionary forces are primarily responsible for constraining the solution space in biological organisms, in which case one should expect consistency across mechanisms used by different members of a species. In other words, different members should use similar (combinations of) solutions. This hypothesis is consistent with our finding of similar results across neural data from the three monkeys in two experiments. However, in other contexts it is also possible that members of a species employ different mechanisms for solving the same task, pointing to the hypothesis that it is the individual learning trajectory that determines the final solution. This hypothesis is supported in experimental data from Pagan et al. (2022), where rats (a different species) were trained on a flexible integration task. Pagan et al. found a diversity of neural mechanisms implemented by different individual rats.

Overall, the question of which factors constrain the solution space may be problem-specific and/or modality-specific. For example, the brain may employ one solution to handle interference between motion stimuli, but leverage a different solution when processing auditory stimuli in a different brain region. In contrast, our results and those of Libby and Buschman suggest a similar solution, namely, the coordinated activity of neurons with both stable and inverting tuning preferences is leveraged for both visual motion and auditory stimuli, and likely generalizes to all sensory modalities and features.

We contrast our approach of exploring an entire solution space with the more common practice of building a single best model to maximally explain neural data. The best-model approach is useful when one is looking to predict a biological system’s response to novel stimuli or any kind of interventions or lesions. However, often the best model has low interpretability and many degrees of freedom to be optimized, which leads this approach to fall short at explaining the underlying mechanisms or principles behind the computations. The solution space approach can uncover the mix of mechanisms behind the computations. It was a deeper understanding of the conceptual solution space that allowed us to engineer

RNNs across a broader range of solutions (Pagan et al., 2022) and ultimately discover those that were most parsimoniously aligned with the brain.

Taken together, our results help elucidate how recurrent neural networks are able to solve the problem of sensory-memory interference by leveraging both static and dynamic codes, and bridges scales from behavior to neural firing patterns to synaptic connectivity. Intriguingly, our work also suggests that, even beyond the specific context of sensory-memory interference, the dynamic neural codes seen in the brain may enable information to effectively “hide” from being overwritten. Finally, we propose a new role for area MST in overcoming sensory-memory interference.

## 4 ACKNOWLEDGMENTS

We thank Laureline Logiaco and Ila Fiete for valuable discussions.

## 5 METHODS

### 5.1 Monkey datasets

The two monkey electrophysiology datasets analyzed in this study are described in Mendoza-Halliday et al. 2014 and Mendoza-Halliday et al. 2024 (Mendoza-Halliday et al., 2014, 2024). In Mendoza-Halliday et al. 2014 monkeys performed a delayed match-to-sample (DMS) task, and in Mendoza-Halliday et al. 2024 monkeys performed a working memory-guided feature attention task. Both experiments required (as a subproblem) overcoming sensory-memory interference, as some trials contained distractor stimuli before the behavioral response. The first experiment used spatially local random dot motion stimuli and the second used full-screen random dot motion stimuli. In the present study, we looked specifically into the neural dynamics during the presentation of the first stimulus and the immediately following delay period. We analyzed 182 and 672 neurons in MST from the studies of Mendoza-Halliday et al. 2014 and 2024, respectively, that were selective for the cue direction in either the cue or delay periods (Figure 6). We analyzed 63 and 64 neurons in MST from the studies of Mendoza-Halliday et al. 2014 and 2024, respectively, that were selective for the cue direction in both the cue and delay periods (Figure 5 and Supplementary Figure S5). Some neurons in the study of Mendoza-Halliday et al. 2024 were recorded simultaneously, which allowed us to compute noise correlations between pairs of simultaneously recorded neurons for repeated presentations of the same stimulus. For a more complete description of the tasks, refer to the original studies (Mendoza-Halliday et al., 2014, 2024).

### 5.2 Recurrent Neural Networks: Architecture and Training

Before discretization, the dynamics of the simulated neurons were governed by the standard continuous-time RNN equation:

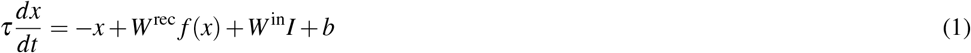

In this study, *τ* =100 ms. The network was simulated using Euler integration with timesteps of *τ/*10 =10 ms. The firing rate of each neuron, *f* =*x*=, was related to its total input *x* through a rectified tanh nonlinearity, *f* =*x*=max=0, tanh=*x*=. All RNNs in this paper contained 100 recurrent units, with the results being largely insensitive to network size. Each of the 100 neurons in the RNN received input from all other neurons through the recurrent weight matrix *W* ^rec^ and also received external input, *I*=*t*=, through the weight matrix *W* ^in^. Firing rates were linearly combined to produce the output *y*=*t*=according to *y* =*W* ^out^ *f* =*x*=. In the training and the analysis phases, Gaussian noise with s.d. 0.1 was added to the firing rate of each neuron at each timestep. All backpropagation-trained RNNs were initialized from a random initialization

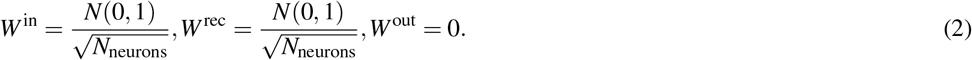

RNNs 5,000 training steps with Adam in Pytorch, with learning rate 0.001 for hand-designed RNNs and 0.0001 for backpropagation-trained RNNs (learning rate had to be lowered for the latter as otherwise they would not converge in training).

#### 5.2.1 Hand-designed RNNs

In the hand-designed networks, entries of *W* ^rec^ and *W* ^in^ were set as *a* cos=*θ*_*i*_ *−θ*_*j*_=, where *θ*_*i*_ stands for the pre-defined preferred direction of neuron *i*. Proportionality constants, the bias *b*, and the linear readout layer were optimized with Adam in Pytorch. For the Gating networks, the recurrent layer was split in two clusters of units (of equal size). The gate units were connected to input and to the ring units using the same cosine connectivity profile, with no recurrent connectivity within the gate units. The ring units were connected recurrently with a cosine connectivity profile, and had additional uniform inhibitory connections back to the gate layer. All proportionality constants were optimized using Adam in Pytorch. For the Strong Attractor networks, the constant a was split into *a*^rec^ and *a*^in^, allowing *a*^rec^ *> a*^in^. For the Reshuffle of Tuning networks, every artificial neuron was assigned a preferred directions for the input and the recurrent connectivity profiles, 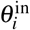 and 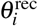 respectively (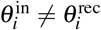 in most cases) 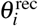 were selected to lie uniformly in the range =0, 360). Then, 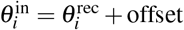. Offsets were selected uniformly at random in the range =*−*130, 130=degrees. The range of offsets was determined by a hyperparameter sweep to produce the network with lowest average error on the task. For the Inversion of Tuning networks, this offset was selected to either be 0° or 180°, with 55% of neurons having an offset of 0° and 45% having an offset of 180°. To create the structural/functional connectivity figure, Strong Attractor and Reshuffle of Tuning solutions were combined as follows: Every artificial neuron was assigned a 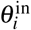 and 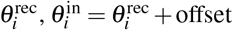, offset was selected uniformly at random in the range =*−A, A*=The proportionality constant a was split into *a*^rec^ and *a*^in^, with *a*^rec^ =*a*^in^ *∗B*. We then considered the whole family of networks for different values of *A* and *B*, where *A* varied from 0° to 160° and *B* varied from 1 to 5. For any given network in the family, the cosine function (with a variable bias and amplitude) was fit to the connectivity profiles, and the extracted amplitude ratios were used to compute the functional and structural ratios.

#### 5.2.2 RNN task

In the Working Memory + Distractor (WM+D) task, the input consisted of a variable delay period (100- 200 ms) with no inputs, then the target cue direction (for 100 ms), followed by a second variable delay (1200-1600 ms). Within this second delay, the distractor direction (sampled randomly and independently of the target direction) was presented, 100-900 ms after the target direction was presented. After the second delay, the network was required to output the target direction for 1000 ms. In the Working Memory Only (WM) task, there was no distractor shown. The task input and distractor (Figure 1b) were modeled with 100 direction-selective neurons, as described by Teich and Qian (2003); the outputs were modeled with sin and cos. In the case of discrete cue directions as in the task of Suzuki and Gottlieb (2013), the target selected by the RNN was taken to be the one nearest the RNN output. During training, all directions (0-360°) were used for the input and distractor.

### 5.3 Data processing

#### 5.3.1 Preprocessing

For further analysis, for every neuron within a network, only its mean firing rate across the cue and delay periods was considered. To exclude the transient responses right after the input stimulus disappeared, the first 200 ms of the delay period was removed in both datasets and in all of the models. For every neuron, a distribution of average firing rates across all experimental conditions was obtained in every window (cue and delay). Then, for every experimental condition, the average firing rate of the neuron in that condition was z-scored in the overall distribution. This z-score was used as the neuron’s normalized firing rate for further analysis. For artificial neurons, smooth z-scoring function was used to avoid dividing by zero in case of no variability in neuronal firing rates:

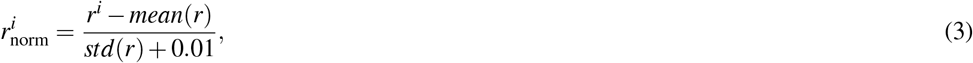

Where 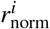 is the normalized (z-scored) firing rate of a given neuron in condition *i, mean*(*r*) and *std*(*r*) are the mean and standard deviation, respectively, of firing rates across all conditions in a given window.

#### 5.3.2 Determining the selectivity of neurons

Feature selectivity (or lack thereof) for any given neuron was determined in the following way. Given the neuron’s window- and trial-averaged firing rates *r*_1_, *r*_2_, …, *r*_*k*_ in response to corresponding stimuli *θ*_1_, *θ*_2_, …, *θ*_*k*_, the Direction Selectivity Index (DSI) was defined as the magnitude of the vector sum of responses, divided by the sum of all responses (Swindale, 1998; Mazurek et al., 2014)

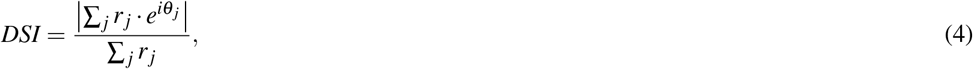

which ranges between 0 (no direction selectivity) and 1 (maximum direction selectivity). The associated preferred direction is the angle of the vector sum:

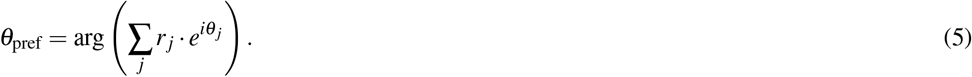

To quantify statistical significance of the selectivity using bootstrap, the analysis above was re-run with all the trial labels reshuffled to obtain the null DSI distribution; the likelihood of the original DSI was tested under that distribution. For the dataset by Mendoza-Halliday et al. 2014 (Mendoza-Halliday et al., 2014), neurons with *p <* 0.01 were considered direction-selective in that time window. For the dataset by Mendoza-Halliday et al. 2024 (Mendoza-Halliday et al., 2014), the threshold was lowered to *p <* 0.005 to correct for higher type 1 error due to many comparisons. We analyzed neurons that were selective in either the cue period *or* the delay period, totaling 182 and 672 neurons from the studies of Mendoza-Halliday et al. 2014 and 2024, respectively. In Supplementary Figure S5 we also computed the Procrustes distance between models and neural responses using a more restricted set of neurons that were selective in both the cue period *and* the delay period, totaling 63 and 64 neurons from the studies of Mendoza-Halliday et al. 2014 and 2024, respectively.

### 5.4 Quantification of distances between networks

To calculate the distance between two networks A and B, their neurons’ window- and trial-averaged responses to four cardinal input stimulus directions (to accommodate the limited available neural data) during the cue and delay task periods were used in the Procrustes distance metric (Ding et al., 2021; Williams et al., 2021). In this study, we used a fully regularized metric with *α* =1.

#### 5.4.1 Distances between networks and neural data

What does it mean for a model to be close to the data? To interpret model-data distance, we need a baseline based on data-to-data distance, which reflects the noise floor present in the data, and is a lower bound that we cannot expect the models to go under. Due to limited subjects, we generated this baseline by splitting the neuronal population in half and comparing the halves to each other, though this may create an overly stringent baseline due to potential neuron dependence.

The data-splitting procedure used for generating the noise floor in Figure 6 is as follows. We split the neural data into nonoverlapping groups each containing *N*_*sample*_ neurons (*ineurons*1, *ineurons*2). We compute the distance between the two samples of neural data *d*1 =*D*=*ineurons*1, *ineurons*2=. *d*1 is the lowest distance we can hope to obtain given the variability in the neurons that were recorded. We sample the same number of *N*_*sample*_ units from the RNN model (*iunits*), and then compute the distance between samples of the model and the neural data *d*2 =*D*=*ineurons*1, *iunits*=. For each iteration of this procedure we get a new estimate for the distance between the model and data, and the data-to-data distance. This procedure was repeated 100 times to obtain representative distributions for each distance.

#### 5.4.2 Low-dimensional embedding

The low-dimensional embedding of the distance matrix was obtained via t-SNE (van der Maaten and Hinton, 2008), using the library openTSNE (Poličar et al., 2024). The perplexity parameter was set to the maximum valid value to maximally preserve the global structure of the data, and the embedding was initialized with the ‘spectral’ initialization.

### 5.5 Code availability

Code for the analyses performed in this study will be made available upon publication.

## 6 SUPPLEMENTARY FIGURES

**Figure S1.**
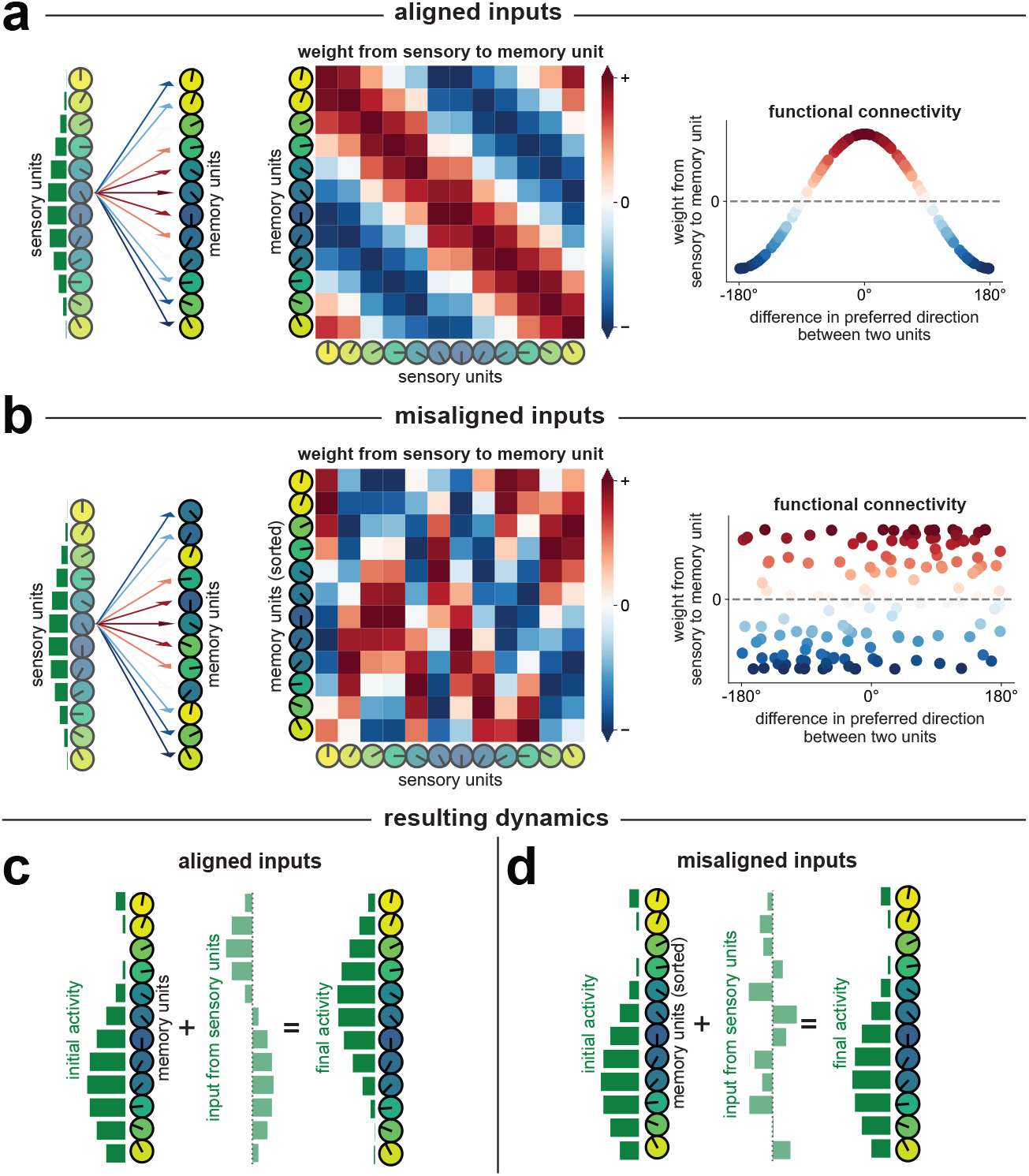
Misalignment of input projections in relation to recurrent connectivity is a strategy for reducing sensory-memory interference. Each circle represents one unit. The color of the circle and orientation of the black line indicates the preferred direction of the unit. In this simplified example there are twelve sensory units and twelve memory units. **(a)** Connections from sensory units to memory units have a cosine connectivity profile with excitation between units with similar preferred directions and inhibition between units with opposite preferred directions. On the left, connections from a single sensory unit to all memory units are shown as colored arrows with red indicating excitatory weights and blue indicating inhibitory weights. In the center, the connectivity from all sensory to memory units is shown as a heatmap. On the right, these same 144 connection weights are plotted as a function of the difference in preferred directions between sensory and memory units, making the cosine connectivity easily visible. A memory unit with a preferred direction of 0 degrees, for example, will receive excitatory inputs from sensory units that have preferred directions near 0. **(b)** The preferred direction of each memory unit changes over time to a new value that is chosen randomly from a uniform distribution centered around the original value. These changes in preferred directions can be caused by, for example, the recurrent connectivity between memory units. As a result of these changes, the weights from sensory to memory units, as a function of the difference in their preferred directions, is no longer coherent (right). A memory unit with a preferred direction of 0 degrees, for example, will now receive both excitatory and inhibitory inputs from sensory units that have preferred directions near 0. **(c)** When the weights between sensory and memory units are aligned, a bump of activity on the sensory units (green bars in panel a) will result in positive inputs to memory units centered around the bump location. This coherent input from the sensory units causes the final activity pattern of the memory units to shift. **(d)** When the weights between the sensory and memory units are misaligned, a bump of activity on the sensory units (green bars in panel b) will result in a mix of both positive and negative inputs to similarly tuned memory units, effectively weakening the sensory input to memory units. The final activity pattern of the memory units will remain largely unchanged.

**Figure S2.**
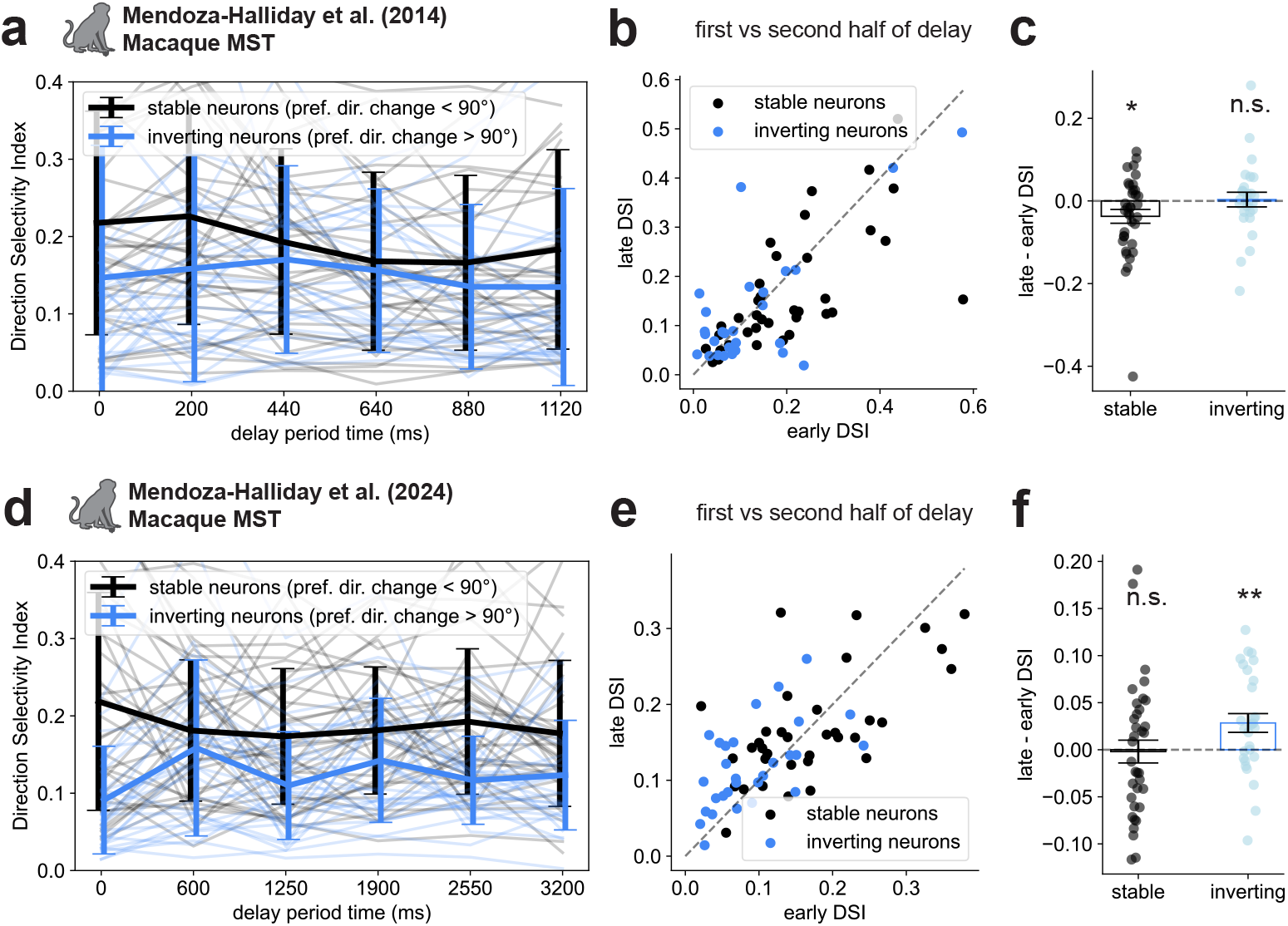
(a, d) The Direction Selectivity Index (DSI) of the stable and inverting neurons does not monotonically decrease during the delay period of the macaque experiments by Mendoza-Halliday et al. 2014 and 2024 (Mendoza-Halliday et al., 2014, 2024). Each line shows the DSI for one neuron over time. Error bars indicate standard deviation across all neurons. (b, e) DSI shown for neural responses averaged across the first versus second half of the delay period. Each dot corresponds to one neuron. (c, f) Differences in DSI from neural responses averaged across the first versus second half of the delay period. Each dot corresponds to one neuron. Same data as in (b, e) except we show the difference in DSI for each neuron. The DSI for the inverting neurons does not decrease during the delay period for both the experiments of Mendoza-Halliday et al. 2014 and 2024. Error bars indicate s.e.m. \**p <* 0.05, \*\**p <* 0.01, Wilcoxon signed-rank test.

**Figure S3.**
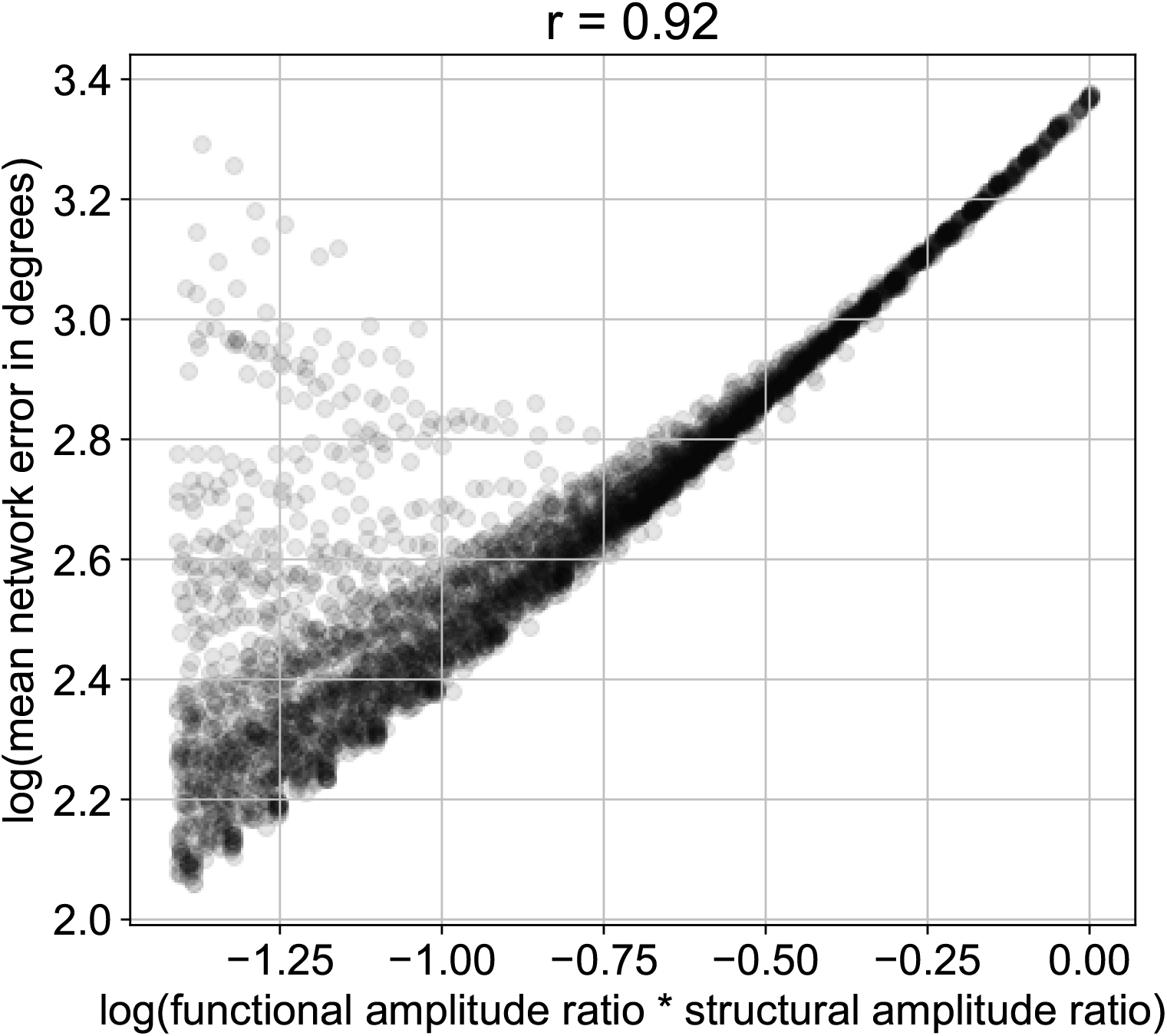
In hand-designed Strong Attractor + Reshuffle of Tuning networks, the product of structural and functional amplitude ratios strongly predicts network performance. N = 4302 networks.

**Figure S4.**
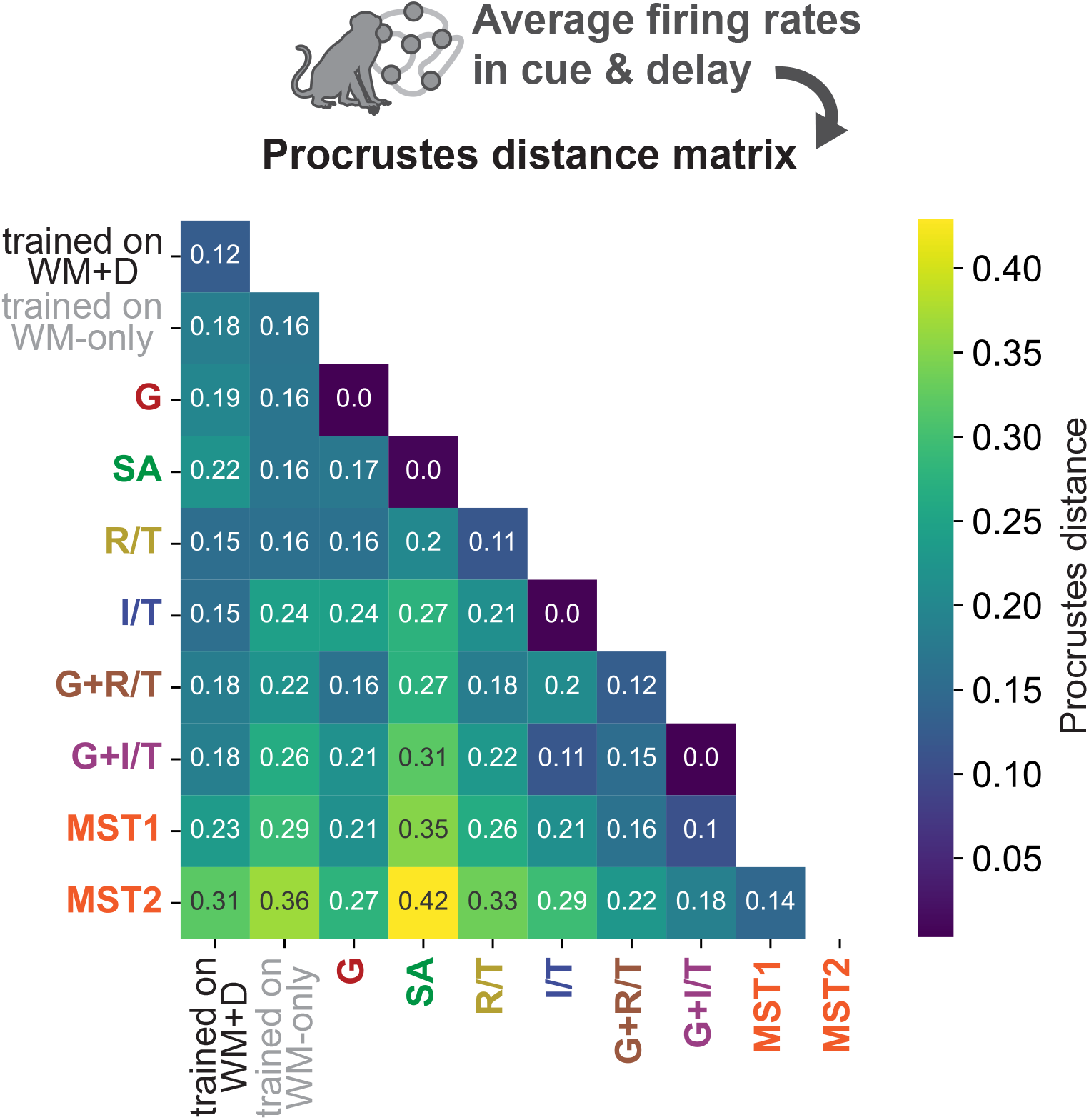
Matrix of pairwise Procrustes distances between each pair of models and neural datasets. A distance of zero indicates perfect alignment. *N* =30 networks per cluster (with networks that did not converge in training omitted).

**Figure S5.**
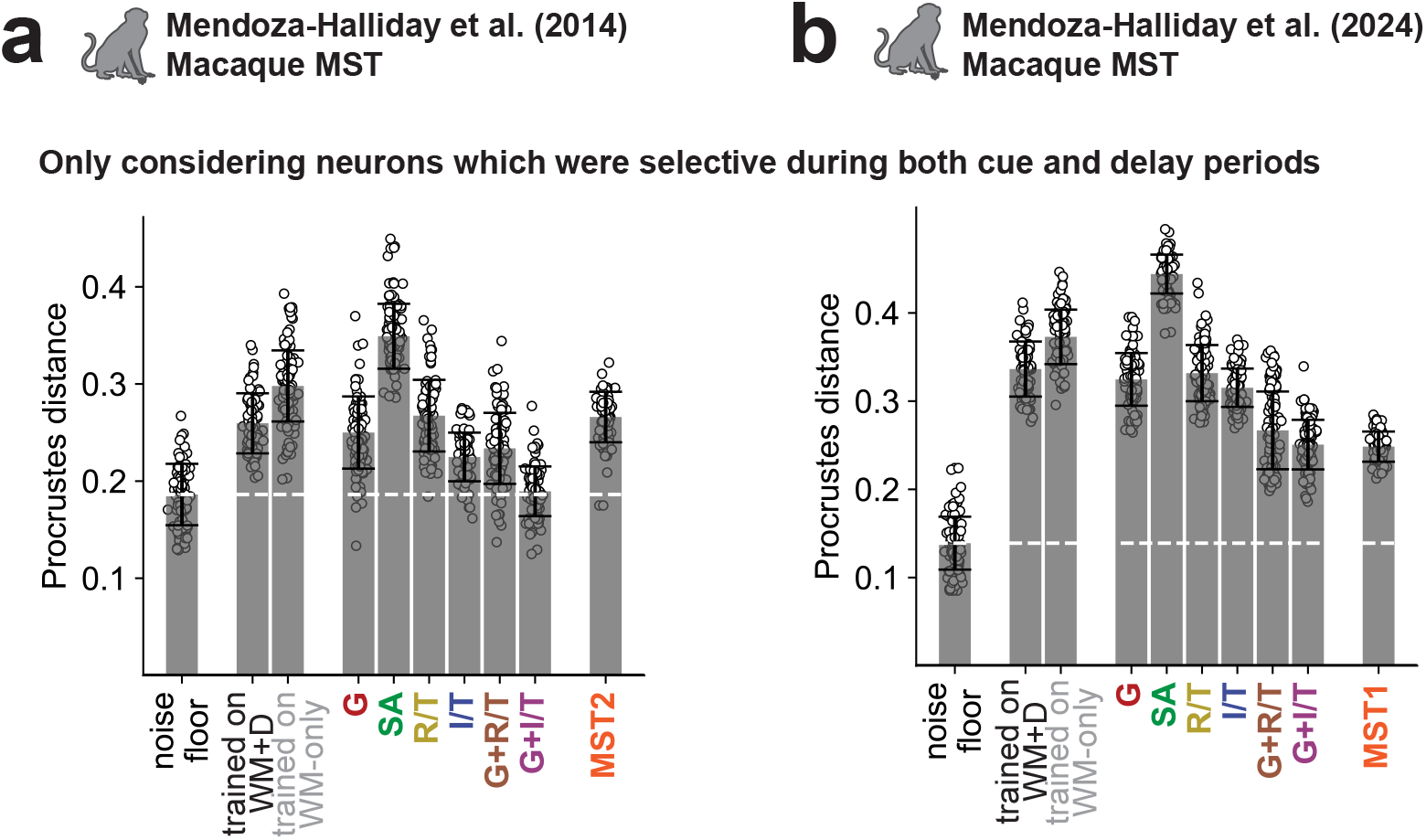
The Gating + Inversion of Tuning mechanism is closest to neural responses in MST irrespective of whether we included neurons selective for the cue direction in both the cue *and* delay periods (above), or in either the cue *or* delay periods (Figures 6b and 6c). Error bars indicate s.d.

**Figure S6.**
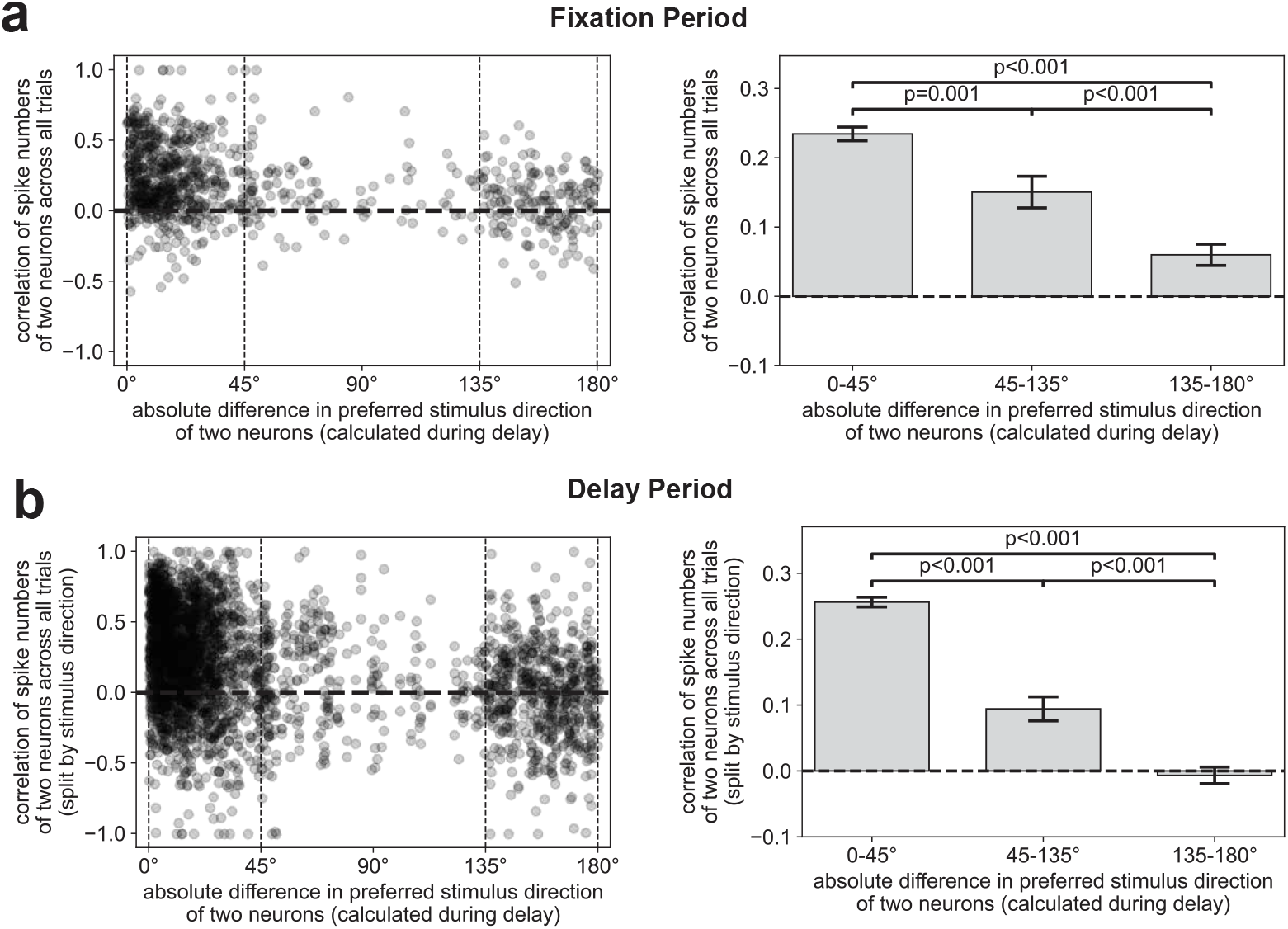
Noise correlations increase for similarly-tuned neurons. Correlations between spike counts of neurons as a function of the absolute difference in preferred stimulus direction, for two task windows: Fixation period (a; *N* =904), and Delay period (b; *N* =3206). Only those neurons that were statistically significantly selective in the delay period are selected. The noise correlations shown here quantify the shared fluctuations between neuron pairs across repeated presentations of the same stimuli. Stimuli were not shown until after the fixation period and so, during the fixation period, all trials were included when computing correlations. Error bars indicate s.e.m

**Figure S7.**
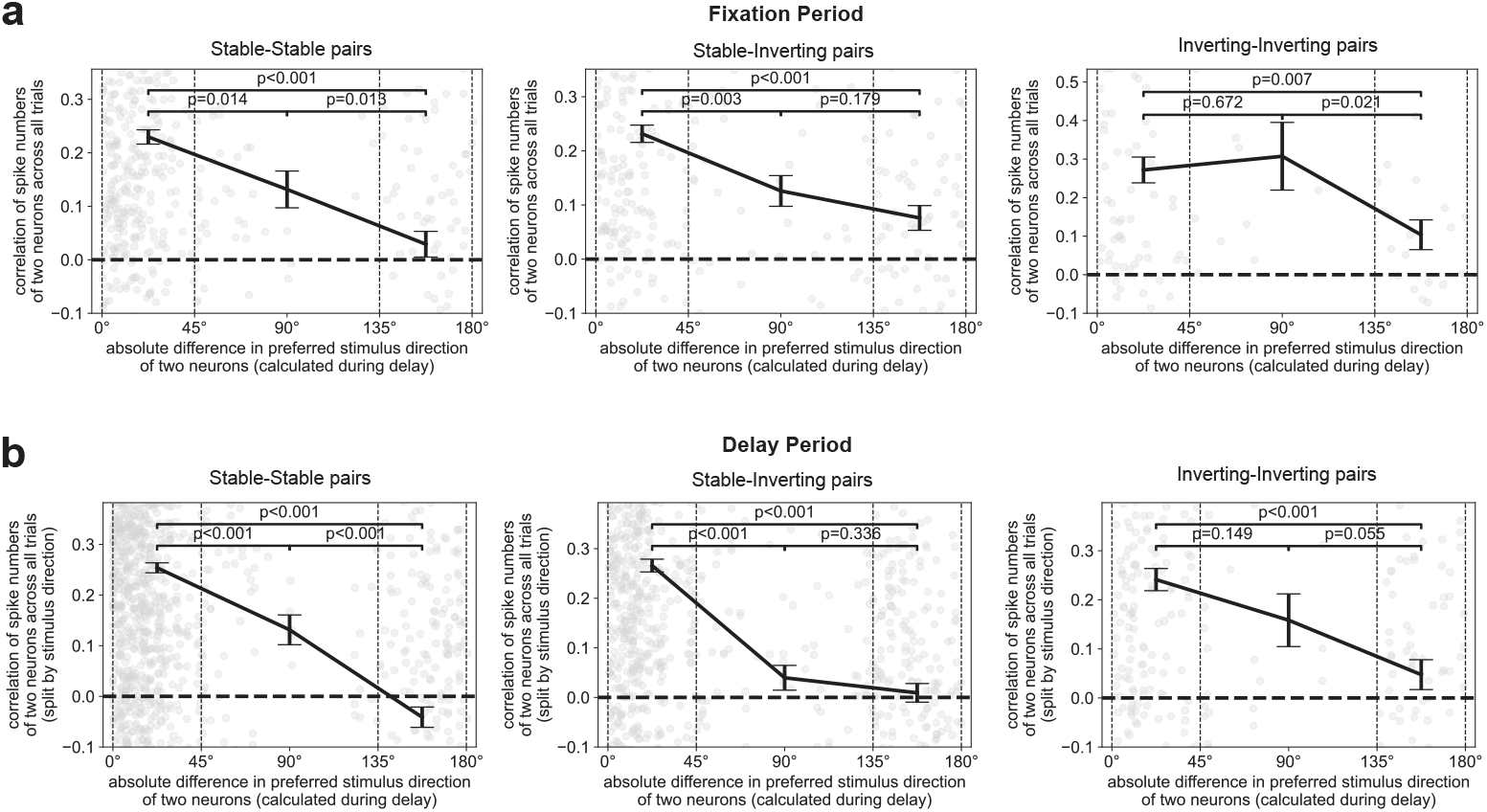
The increase in the noise correlations for similarly-tuned neurons is present within and across stable and inverting clusters of neurons. For all panels: left, stable-stable pairs (preference change *<* 90° for both neurons); middle, stable-inverting pairs (preference change *<* 90° for one neuron and *>* 90° for the other); and inverting-inverting pairs (preference change *>* 90° for both neurons). Only those neurons that were statistically significantly selective in the delay period are selected. Task windows: Fixation period (a; *N*_stable_ =493, *N*_mixed_ =310, *N*_inverting_ =101), and Delay period (b; *N*_stable_ =1726, *N*_mixed_ =1120, *N*_inverting_ =360). The noise correlations shown here quantify the shared fluctuations between neuron pairs across repeated presentations of the same stimuli. Stimuli were not shown until after the fixation period and so, during the fixation period, all trials were included when computing correlations. Error bars indicate s.e.m

## Notes

### Competing Interest Statement

The authors have declared no competing interest.

### Summary of Updates

Added figure and text on neural adaptation.

